# MATR3 is essential for oocyte growth and maturation quality through a dual molecular mechanism

**DOI:** 10.64898/2026.02.12.705648

**Authors:** Yibing Bao, Zhenzi Zuo, Tengteng Wang, Lin Lin, Meng Gao, Shaogang Qin, Qingfeng Yang, Bingying Liu, Wanyuan Sun, Jie Ma, Tianhua Zhu, Guoliang Xia, Bo Zhou, Rong Hu, Hua Zhang, Fengchao Wang, Chao Wang

## Abstract

The molecular mechanisms governing mRNA accumulation during oocyte growth, essential for developmental competence, remain poorly understood. This study investigates the role of Matrin-3 (MATR3), a highly expressed RNA-binding protein in growing oocytes (GOs), using oocyte-specific knockout mouse models and human oocyte maturation arrest (OMA) samples. The results showed that MATR3 was more abundant in GOs than fully-grown oocytes (FGOs), highly expressed in the nucleus of non-surrounded nucleolus (NSN) oocytes, and exited the nucleus during the NSN-to-surrounded nucleolus (SN) transition. In OMA patients, MATR3 nuclear localization was missed, with smaller oocytes than FGOs. Further, *Matr3* deletion in mouse GOs caused restricted oocyte growth, global transcription disorders, follicle development failure, blocked GO-granulosa cell communication (via reduced *Gdf9* and *Radixin* expression), and infertility. Mechanistically, MATR3 regulated transcription by recruiting H3K9me2-demethylating lysine-specific demethylase 3B or binding target gene promoters, like *Radixin*. These findings reveal a critical role of MATR3 in orchestrating transcription and paracrine signaling during oogenesis and suggest its potential as a diagnostic and therapeutic target for OMA.

## Introduction

The growth of oocytes refers to the process in which growing oocytes (GOs) develop into fully grown oocytes (FGOs). Oocyte growth involves an increase in volume, as well as the accumulation of metabolic molecules, transcripts and proteins. *In vivo*, FGO determines the developmental potential of early embryo. A high qualified FGO derives from the GO in a growing follicle, which may experience dynamic gene transcription as well as finetuned orchestration between the GO and the surrounding granulosa cells. In the clinic, besides genetic caused female infertility, uncovering the etiologies especially epigenetic causative reasons is equally pivotal for improving the oocyte maturation quality *in vitro* (Ren et al. 2017; Wang et al. 2019; He et al. 2021; Gao et al. 2024; Zhu et al. 2024). Patients who suffer from oocyte maturation arrest (OMA) are unable to retrieve mature oocytes during their repetitive *in vitro* fertilization (IVF) and intracytoplasmic sperm injection (ICSI) cycles (Liu et al. 2018). In the past, the importance of oocyte specific RNA binding proteins (RBPs) like MARF1, LSM14B, and MTR4 in improving oocyte development as well as their relationship to OMA have been confirmed (Su et al. 2012; Su et al. 2021; Wu et al. 2025). However, it remains unclear how other RBPs coordinates epigenetic molecules contributing to transcribe and preserve massive mRNAs to direct the timely growth of GOs and growing follicles.

Multiple RBPs have been proven to closely correlate to OMA. The PATL2 protein is known to regulate the homeostasis of maternal mRNAs, while a whole exome sequencing study has identified four recessive variants of the *PATL2* gene in three OMA families (Wang et al. 2024a). Also, the compound heterozygous *ZFP36L2* mutations correlates to OMA (Wan et al. 2024). ZFP36L2 is crucial for regulating the degradation of maternal mRNAs and the maturation of mouse oocytes (Chousal et al. 2018). Despite of these, it is unclear if other RBPs contribute to female fertility as well. This study focuses on a conserved RBP Matrin-3 (MATR3) who contains two zinc-finger domains to specifically bind to DNA sequences rich in GC (Zeitz et al. 2009). The two RNA recognition motifs of MATR3 participates in the regulation of alternative splicing by specifically binding to mRNA sequences rich in AU (Zeitz et al. 2009; Iradi et al. 2018). Moreover, MATR3 is rich in nuclear localization sequences, which enabling it to localize in the nucleus. One of the characteristic functions of MATR3 is that it recruits other RBPs while binding to the target sequences (Pollini et al. 2021). Importantly, as a powerful RBPs, the abnormal aggregation and pathological mutations of MATR3 may interfere with normal cellular functions, including RNA stability and transport (Coelho et al. 2015; Cha et al. 2021). Unfortunately, whether and how MATR3 correlates to GO to FGO development and female fertility remains unknown.

The growth of growing follicles as well as oocyte maturation depends strictly on endocrine and paracrine regulations *in vivo*. Studies including ours have shown that during cyclic recruitment of growing follicles in each estrus cycle, paracrine factor like CNP (C-Type Natriuretic Peptide) coordinates gonadotropins to timely control FGO meiosis arrest, meiosis resumption and ovulation (Zhang et al. 2010). However, throughout the progress from GO to FGO, cumulative studies have highlighted the critic roles of oocyte-secreted factors (OSFs) in supporting follicular somatic cell proliferation, metabolism and GO development (Hanrahan et al. 2004; Su et al. 2008; Nicol et al. 2009; Silva et al. 2011; Gao et al. 2024). Of which, TGFβ superfamily members growth differentiation factor 9 (GDF9) and bone morphogenic protein 15 (BMP15) are the most studied ones (Dong et al. 1996; Dube et al. 1998). Within a growing follicle, GDF9 functions by regulating the cuboidal transformation and promoting proliferation of granulosa cells, which has been repetitively demonstrated by studies including ours (Carabatsos et al. 1998; Spicer et al. 2008; Gao et al. 2024). BMP15, a highly homologous to GDF9, regulates the development of early growing follicles by controlling the proliferation of granulosa cells through the regulation of the GDF9-SMAD signaling pathway (Gilchrist et al. 2008). Most recently, we have proved that the major function of oocyte-derived mushroom-like microvilli (Oo-Mvi) relates to the release of OSFs timely and orderly. Loss of RADIXIN (RDX), the key component of Oo-Mvi, resulted in a failure of the formation of Oo-Mvi and a shortened reproductive lifespan in females (Zhang et al. 2021). However, how RBPs regulates OSFs like GDF9 expression and secretion via Oo-Mvi remains unsure.

Results of this study showed that MATR3 is highly expressed in the GOs of mice, pigs, and humans, indicating a conserved and essential function. MATR3 is required for female fertility, as its specific deletion in mouse GOs severely reduced antral follicle formation and resulted in infertility. MATR3 regulates transcription by recruiting H3K9me2-demethylating lysine-specific demethylase 3B or binding target gene promoters, like *Radixin*. Conclusively, MATR3 is a strong candidate causative RBP contributing to OMA in females.

## Results

### 1. MATR3 expression in oocyte of growing follicle is required for female fertility

To investigate the potential function of MATR3 in female reproduction, we examined its location and expression pattern during folliculogenesis in multiple mammals. The results showed that MATR3 was generally expressed in the nucleus of oocytes and somatic cells of mice (Fig.1A, Fig.S1A). Specially, the level of MATR3 in GOs remained at relatively higher levels than those found in the FGOs as well as stages after maturation (Fig.1B, C), implying that MATR3 may be more important for supporting mouse GO growth. Interestingly, compared to the normal group in which the expression pattern of MATR3 is similar to that of mouse, we noticed a decrease in MATR3 protein level in FGO (NSN) from a patient with OMA, which has not been reported before (Fig.1D). This finding suggests that MATR3 may be one of a candidate responsible for OMA. Similarly, we also detected the expression pattern of MATR3 in the ovaries of porcine, which is highly consistent with that in mice and humans, indicating that the physiological function of MATR3 may be highly conserved among species (Fig.S1B-D).

**Fig 1.**
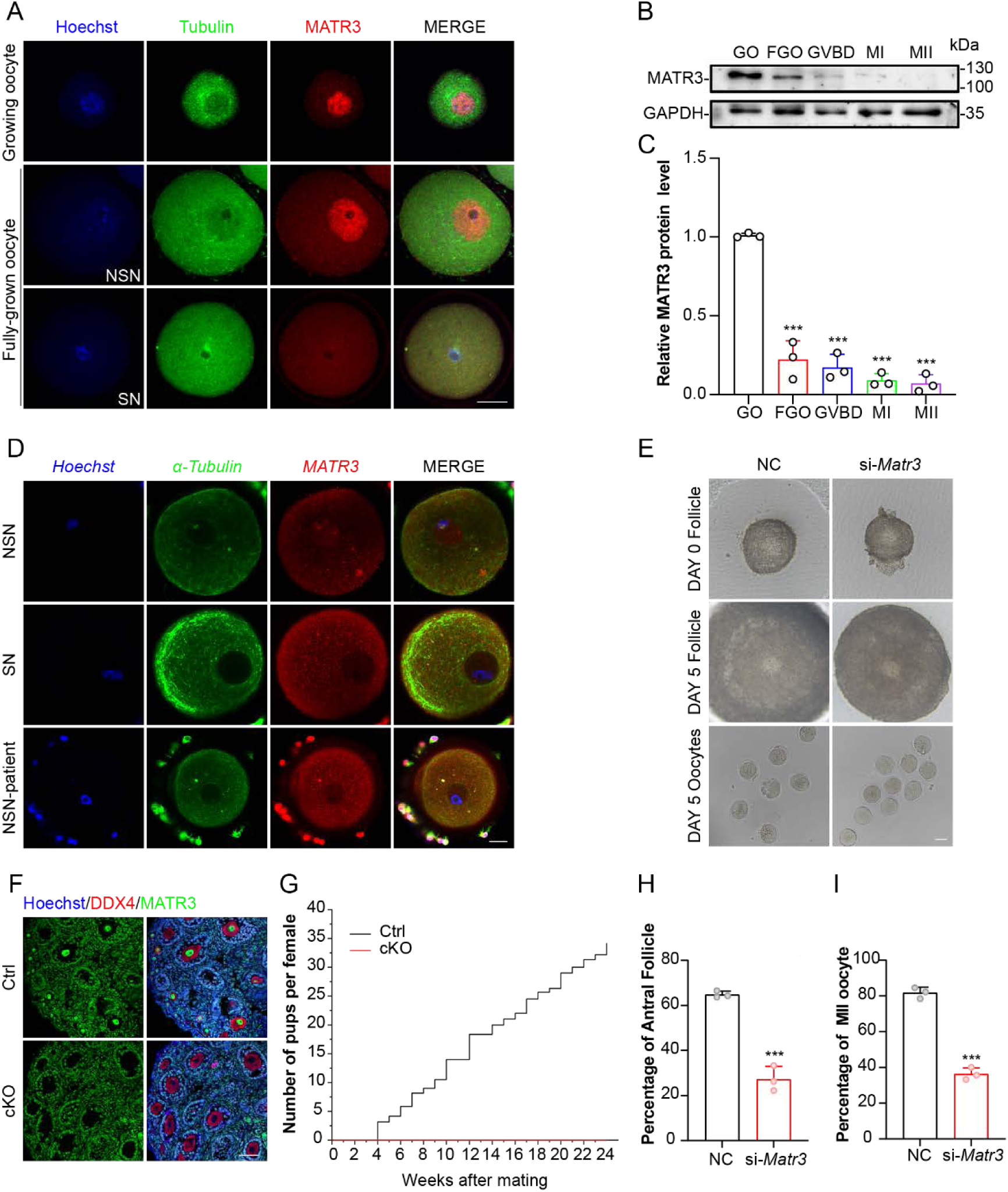
MATR3 in growing oocyte regulates female reproduction in mice. **A** Immunofluorescence showing MATR3 levels in mouse oocytes. **B** Western blot results showing MATR3 expression in mouse oocytes. Total proteins from 200 oocytes were loaded in each lane. GAPDH served as a loading control. **C** Relative expression level of MATR3 to β-Actin. **D** Immunofluorescence showing MATR3 levels in human oocytes. n = 11. **E** Representative image of follicles and oocytes from NC and si-*Matr3.* **F** Immunohistochemistry results showing MATR3 expression in oocytes. **G** Cumulative number of pups from Ctrl (n = 6) and cKO (n = 3) female. **H** Percentage of antral follicles in **e**. n ≥ 33 per group. **I** MII ratio of oocytes collected from **e**. n ≥ 33 per group. Scale bars: 20 μm in **A**, **B** and **C**, 40 μm in **E** and **F**. Data are represented as mean±SD. \*\*\**P* < 0.001, \*\**P* < 0.01, \**P* < 0.05, n.s., not significant.

To explore the effect of MATR3 on GO growth, the authors firstly established an oocyte knockdown model of early growing follicles in mouse. This model enables the development of secondary follicles (SFs) into antral follicles (AFs) and, successful ovulation after gonadotropins hormone supplementation (Fig.S2A). After injecting *Matr3* siRNA into the oocyte of SF with a diameter of 150 μm in the model, these follicles were *in vitro* cultured for 5 - 7 days (Fig.S2B-D). Photographs were taken daily to record the follicular development status. On the fifth day, oocytes and granulosa cells were collected respectively to detect the extrusion rate of the first polar body of oocytes and multiple functional markers of granulosa cells. The results showed that after knocking down *Matr3* in the oocytes of SFs, follicular development was hindered. Specifically, when cultured *in vitro* for five days, the SFs in the negative control group (NC) could develop into AFs, while the proportion of AFs in the knockdown group (si-*Matr3*) was significantly reduced (Fig.1E, H 64.87±1.531% vs 27.27±5.613%). Consistently, the extrusion rate of the first polar body of oocytes was significantly reduced (Fig.1I 81.83±3.048% vs 36.35±3.380%). Meanwhile, the protein level of FOXL2 in granulosa cells decreased, the protein level of the proliferation marker PCNA decreased, and the protein level of the hormone-responsive receptor IGFIR decreased as well (Fig.S2E).

Based on the above *in vitro* experiments, we utilized a novel mouse model with specific MATR3 deficiency in oocytes to explore the MATR3 function. Specifically, *Matr3* was conditionally knocked out after 3 dpp using *Gdf9*-Cre mice to generate *Matr3*^flox/flox^; *Gdf9*-Cre (cKO) mice and to evaluate its specific effect in oocytes (Fig.S3A). The knockout efficiency of cKO mice was examined by immunofluorescence (Fig.1F, Fig.S3B). Notably, a fertility test showed that cKO mice were infertile during the 6-month mating process (Fig.1G). Thus, MATR3 in oocytes is crucial for the growth of follicles and the maintenance of fertility in mice.

### 2. MATR3 is indispensable for supporting GO growth

To clarify how MATR3 deficiency hindered mouse fertility, we injected human chorionic gonadotropin (hCG) after intraperitoneal injection of pregnant mare serum gonadotropin (PMSG) for 48 h to the mouse at postnatal day 23 (PD23), and isolated the ovaries and oocytes after treated for 13 h (H13). We evaluated the number of follicles at all developmental stages in the ovaries of H13. The results showed that there were no corpora lutea (CL) in the ovaries of cKO mice (Fig.2A, B). Consistently, hCG failed to induce ovulation in cKO mice (Fig.2C, D). Meanwhile, both the number of AFs (Fig.2B) and the ratio of AFs/SFs (Fig.2L) were significantly lower in cKO mice than in the control (Ctrl). GOs from PD14 cKO mice had normal size and morphology (Fig.S3C, D). While, FGOs from PD23 cKO mice had smaller sizes than Ctrl oocytes (Fig.2E, F) and the ratio of SN-type oocytes in cKO was significantly lower than that in the Ctrl group (Fig.2G). During *in vitro* maturation, the proportion of oocytes from PD23 cKO mice developing to metaphase II stage was significantly reduced (Fig.2E, H 54.9±2.08% vs 9.57±1.11%).

**Fig 2.**
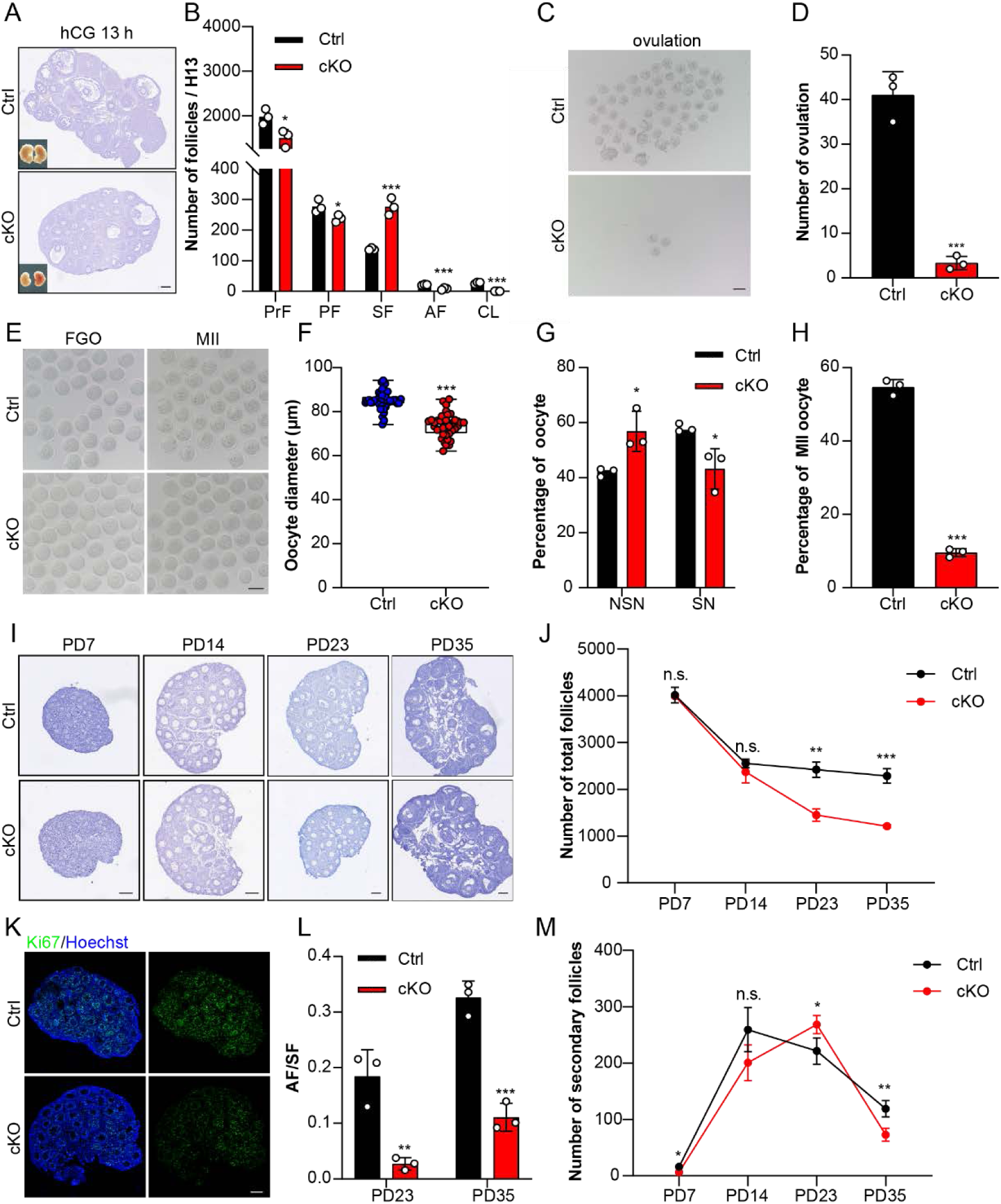
MATR3 was required to support oocyte growth. **A** Hematoxylin staining of PD23 ovaries after gonadotropins injection. **B** Number of follicles in PD23 ovaries after gonadotropins injection. n = 3. **C** Morphology and number of oocytes collected from oviducts of PD23 mice after superovulation. n = 3. **D** Morphology of oocytes derived from PD23 mice primed with PMSG for 48 h. n = 3. **E and F** Diameter of oocytes derived from PD23 mice primed with PMSG for 48 h. n = 40. **G** NSN and SN ratio of oocytes derived from PD23 mice primed with PMSG for 48 h. n = 3. **H** MII ratio of oocytes derived from PD23 mice primed with PMSG for 48 h. n = 3. **I** Hematoxylin staining of PD7, PD14, PD23, PD35 ovaries. N =3. **J** Number of total follicles in PD7, PD14, PD23, PD35 ovaries. n = 3. **K** Cell proliferation indicated by Ki67-positive GCs in mice ovaries. **L** Statistics of AF/SF in PD23 and PD35 ovaries. n = 3. **M** Number of SFs in PD7, PD14, PD23, PD35 ovaries. n = 3. Scale bars: 40 μm in **C** and **E**, 120 μm in **A**, **I** and **K**. Data are represented as mean±SD. \*\*\**P* < 0.001, \*\**P* < 0.01, \**P* < 0.05, n.s., not significant.

To investigate the inner mechanisms of female infertility in cKO mice, we firstly detected the development of cKO ovaries at different ages (Fig.2I, J). At PD7, a similar morphology of ovaries was found in cKO and Ctrl mice, showing that oocyte deletion of *Matr3* does not affect the formation or survival of dormant primordial follicles (Fig.S3E). However, the total follicles began to lost at PD14, and a significant difference began to appear at PD23. This was reinforced by the data of available follicles within the ovaries of mice at PD35 (Fig.2I, J). Further statistical results showed that the development of SFs was slowed down from PD7 to PD23, and the conversion rate to AFs was significantly reduced at PD23 (Fig.2M). Consistently, immunofluorescence staining showed that the numbers of Ki67-positive (Fig.2K, Fig.S9) in cKO mice were lower than those found in the Ctrl (73.55±13.29% vs 24.65±7.80%).

In summary, the deletion of *Matr3* lead to impaired growth of oocytes. Oocyte with impaired growth induced a follicle development disorder in a time-dependent manner attributed to the arrested transition of secondary-to-antral follicles.

### 3. MATR3 governed global transcription in GO is pivotal for the growth of follicle

To investigate the differences of transcription levels in oocytes between the GOs and FGOs, oocytes were isolated based on their diameters. During the time, 5-ethynyl uridine (EU) was used as probes to measure the levels of newly synthesized mRNA in cells. The results indicated a potential role for MATR3 in the maintenance of high transcriptional level of GOs, which showed that he expression of MATR3 was consistent with the transcriptional activity of oocytes (Fig.S4).

To identify the genes regulated by MATR3, the NC and si-*Matr3* oocytes were used for single-cell RNA-sequencing (scRNA-seq). The results of PC clustering analysis and heat map showed good data parallelism (Fig.S5). The results showed that 1552 genes were upregulated and 1155 genes downregulated, respectively (|Foldchange|≥2, *P*value<0.05, Fig.3A). Gene ontology (GO) enrichment analysis showed that the DEGs were enriched in the terms related to fibrillar centers, chromatin organization, and nucleotide binding (Fig.3B).

**Fig 3.**
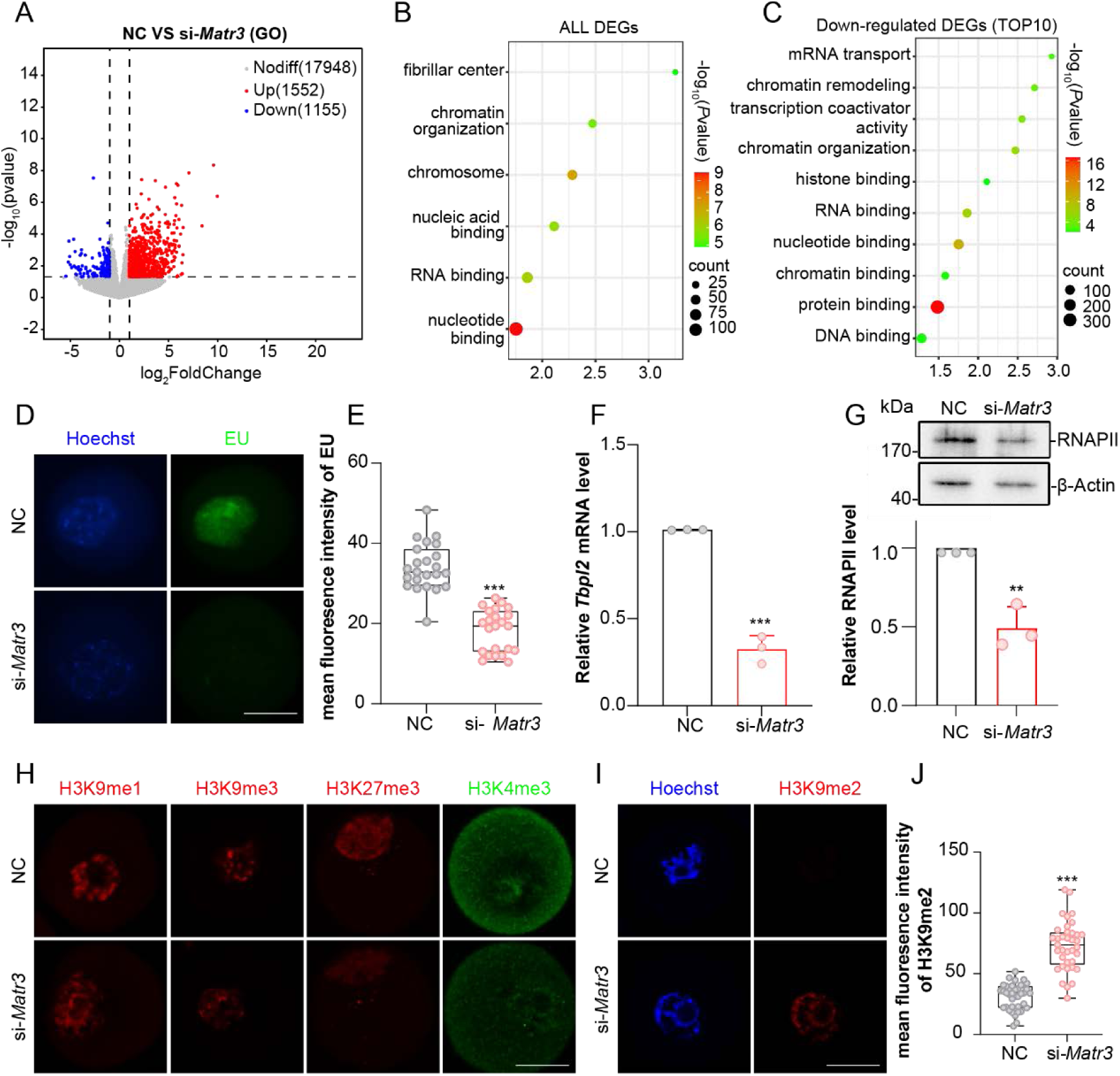
*Matr3* knockdown results in the reduction of transcriptional activity. **A** Volcano plot shows DEGs (upregulated, red; downregulated, blue) in si-*Matr3* oocytes compared to the NC. **B** Key GO enrichment of all DEGs in GOs. **C** TOP10 terms of downregulated DEGs in GOs. **D** EU staining (green) in GO collected from NC and si-*Matr3.* n ≥20. **E** Quantification of the mean fluorescence intensity of EU in oocytes. **F** RT-qPCR results showing *Tbpl2* mRNA level in GOs. **G** Western blotting results showing RNAPII protein level in GOs. **H** H3K9me1 (red), H3K9me3 (red), H3K27me3 (red), H3K4me3 (green), staining in GO collected from NC and si-*Matr3.* n ≥20. **I** H3K9me2 staining (red) in GO collected from NC and si-*Matr3.* n ≥37. **J** Quantification of the mean fluorescence intensity of H3K9me2 in oocytes. Scale bar: 20 μm. Data are represented as mean±S.D. \*\*\**P* < 0.001, \*\**P* < 0.01.

Subsequently, we performed GO analysis on the down-regulated differentially expressed genes in oocytes. We found that approximately two-thirds of the top 10 entries were related to transcription (Fig.3C). Based on this, we performed EU staining on oocytes to detect the level of their newly synthesized mRNA. The results showed that, after *Matr3* knockdown, the transcriptional activity of GOs was significantly reduced (Fig.3D, E). The relative mRNA level of the transcription initiation factor *Tbpl2* (Yu et al. 2020), which is specifically expressed in oocytes, was significantly decreased (Fig.3F). Moreover, the relative protein level of RNAPII fell significantly (Fig.3G).

Through GO enrichment analysis of scRNA-seq, it showed that the differentially expressed genes were significantly enriched in the “chromatin organization” term, which was highly related to transcription. To prove that MATR3 affects the transcriptional activity of oocytes by regulating the chromatin state, we used immunofluorescence to detect the methylation levels of multiple histones, aiming to find the key molecules responsible for the downregulation of oocyte transcriptional activity. After knocking down *Matr3* in GOs, the protein levels of H3K9me1 and H3K9me3, which were related to gene transcriptional repression, did not change significantly. Contrarily, the protein levels of H3K27me3 and H3K4me3 even decreased significantly (Fig.3H, Fig.S6). The above results indicate that H3K9me1, H3K9me3, H3K27me3, and H3K4me3 do not play a dominant role in the regulation of GO transcriptional levels mediated by MATR3. Excitingly, the protein level of H3K9me2, which is related to gene transcriptional repression, increased extremely high. The level of H3K9me2, which marks transcriptional repression, was significantly increased either (Fig.3I, J). Based on the above results, MATR3 in GO maintains a high transcriptional level by eliminating the inhibitory modification H3K9me2.

### 4. MATR3 governs follicle growth partially through regulating GDF9 production

During the follicle growth process, a high transcriptional level in GO is a prominent physiological feature. The substantial accumulation of maternal mRNAs, proteins, and metabolites in the cytoplasm of GO is a crucial factor contributing to the gradual increase in oocyte volume. It is also essential for the development of early-growing follicles. Among these, GDF9 plays a vital role in regulating the development of early-growing follicles. Specifically, we examined the key factors related to the GDF9 signaling pathway to verify the paracrine signals.

The results showed that the *Gdf9* level was decreased significantly in cKO ovaries, which was confirmed by the mRNA and protein expression assays (Fig.4A-C). When *Matr3* in oocytes was knocked down *in vitro*, the level of GDF9 was significantly decreased consistently (Fig.4E, F). Generally, the GDF9 signal is transmitted to SMAD3 in the cytoplasm of granulosa cells and enters the nucleus through phosphorylation, thereby promoting the differentiation of granulosa cells. Compared with the Ctrl group, the level of phosphorylated SMAD3 protein was decreased in cKO ovaries (Fig.4B). Although there was no significant difference in SMAD3 protein levels, most of SMAD3 molecules could not be phosphorylated and was therefore located in the cytoplasm rather than inside the nucleus of the granulosa cells (Fig.4D).

**Fig 4.**
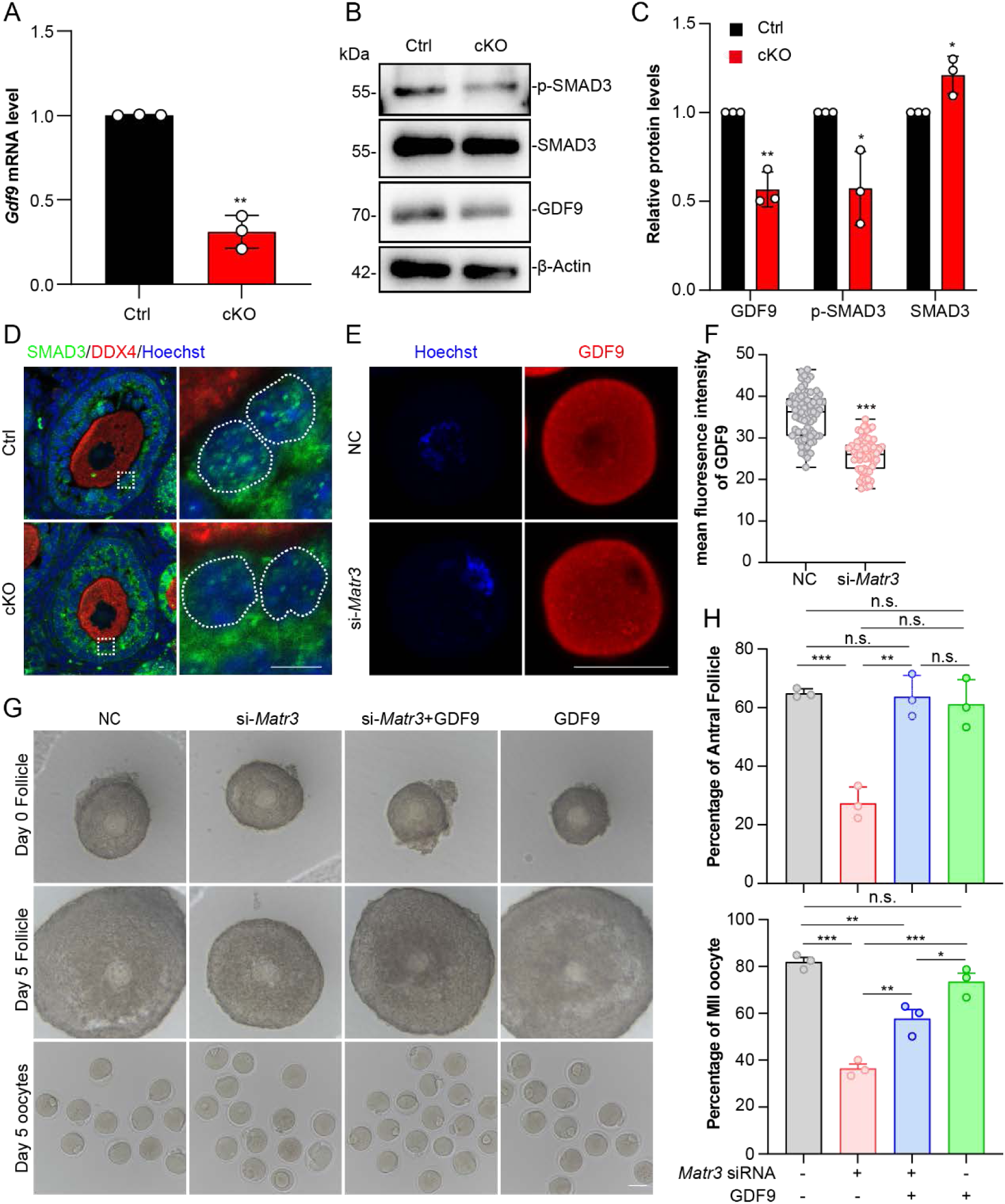
MATR3 controls oocyte growth by regulating GDF9 level. **A** RT-qPCR results showing *Gdf9* mRNA level in PD14 ovaries. **B and C** Western blotting results showing GDF9, SMAD3 and p-SMAD3 protein levels in PD14 ovaries. **D** p-SMAD3 staining (green) in PD14 ovaries. **E** GDF9 staining (red) in GOs from NC and si-*Matr3*. n ≥68. **F** Quantification of the mean fluorescence intensity of GDF9 in oocytes. **G** Representative image of follicles and oocytes from NC, si-*Matr3*, si-*Matr3*+GDF9, GDF9. **H** Percentage of antral follicles and MII oocytes in **G**. n≥3 per group. Scale bar: 40 μm in **D** and **E**, 80 μm in **G**. Data are represented as mean±S.D. \*\*\**P* < 0.001, \*\**P* < 0.01, \**P* < 0.05, n.s., not significant.

To prove that GDF9 is a key downstream molecule whose production is affected by MATR3, we conducted a rescue experiment on growing follicles with *Matr3*-knocked-down oocytes by adding pure GDF9 *in vitro*. We divided the *in vitro* cultured follicles into four groups: the negative control group, the *Matr3*-knocked-down group, the group with GDF9 added after *Matr3* knockdown, and the group with GDF9 added alone. Consequently, after knocking down *Matr3* in the oocytes of SFs and culturing them *in vitro* for 5 days, we photographed and recorded the follicle diameters, and counted the extrusion rate of the first polar body of oocytes (Fig.4G). The results showed that the follicle diameter of the group with GDF9 added after *Matr3* knockdown significantly rescued that of the group with *Matr3* knocked down alone. And, there was no significant difference compared with the control group. The extrusion rate of the first polar body in the group with GDF9 added after *Matr3* knockdown also significantly rescued that of the group with *Matr3* knocked down alone, but there was still a significant difference compared with the extrusion rate of the first polar body in the NC group (Fig.4H). These results indicate that GDF9 can partially rescue the follicular development arrest caused by *Matr3* knockdown and is a key functional molecule in MATR3 - regulated follicular development.

### 5. MATR3 promotes the expression of *Gdf9* by recruiting KDM3B

To explore how MATR3 is involved in the regulatory synthesis of GDF9, we detected the level of lysine (K)-specific demethylase 3B (KDM3B), which directly regulates H3K9me2. Under physiological conditions, KDM3B is localized in the cytoplasm and nucleus of oocytes, and is highly expressed in GO (Fig.S7). This study showed that KDM3B was downregulated as the transcriptional activity of oocytes decreases (Fig.5A, B). To further clarify how MATR3 regulates KDM3B, previous literature indicates that RBPs can recruit KDM3B to regulate the level of H3K9me2 (Kim et al. 2012; Li et al. 2020). Then, to prove this hypothesis, we detected the interaction between MATR3 and KDM3B in the HEK293T-cell line using co-immunoprecipitation (CO-IP). The results showed that there was an interaction between MATR3 and KDM3B (Fig.5C, D). To identify the specific domains of MATR3 for recruiting KDM3B, we firstly constructed the MATR3-EGFP fusion protein. The Western blotting results demonstrated the successful construction of the vector after we transfected the HEK293T-cell line with the constructed plasmids (Fig.S8A, B).

**Fig 5.**
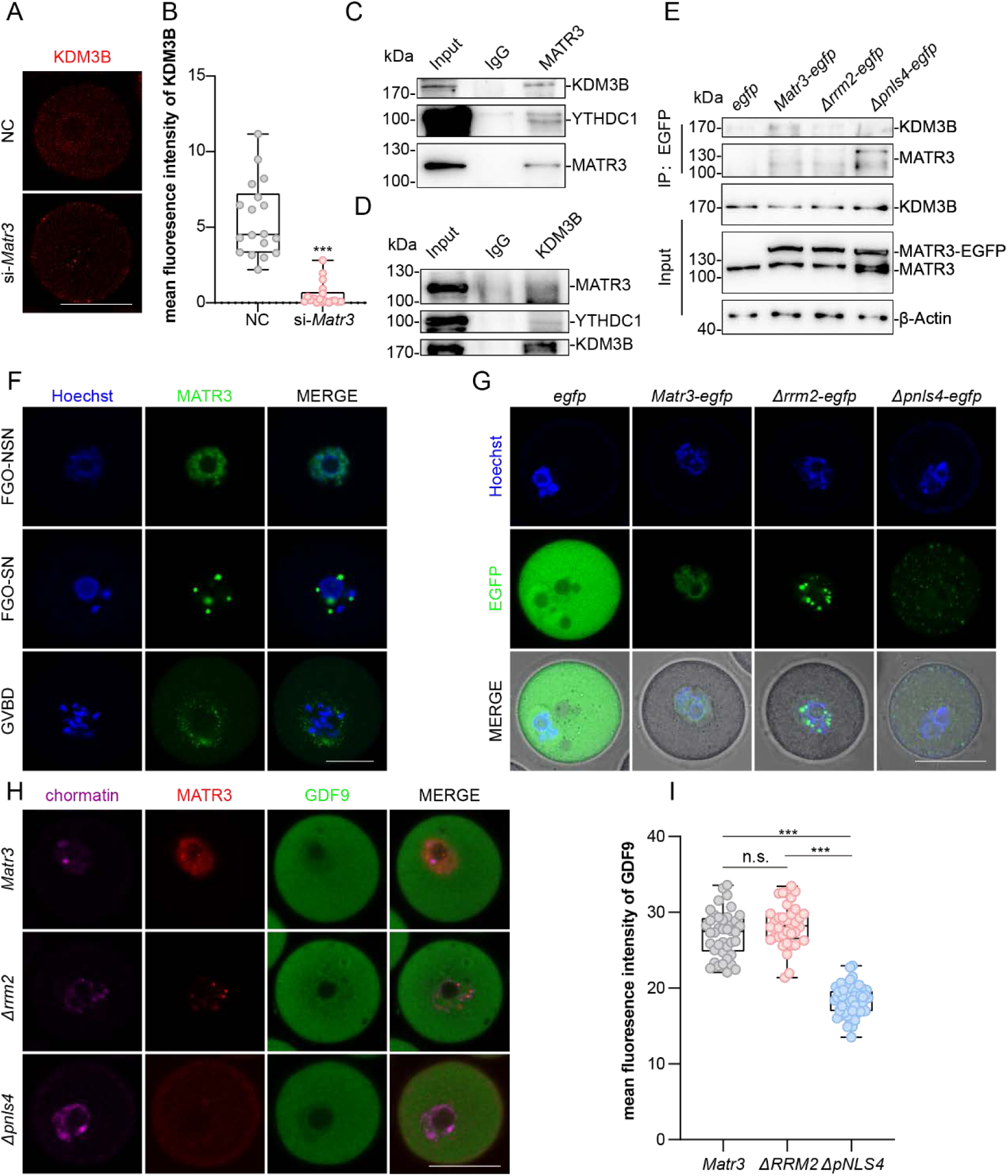
MATR3 promoted GDF9 by recruiting KDM3B. **A** KDM3B staining (red) in GO collected from NC and si-*Matr3*. n ≥18. **B** Quantification of the mean fluorescence intensity of KDM3B (**A**) in oocytes. **C and D** CO-IP showing protein interactions between MATR3 and KDM3B in HEK293T cells. **E** CO-IP results after HEK293T cells were treated with pCDNA3.1-EGFP, pCDNA3.1-MATR3-EGFP, pCDNA3.1-ΔRRM2-EGFP or pCDNA3.1-ΔpNLS4-EGFP plasmid. **F and G** Live-cell imaging of oocytes after injecting *Matr3-egfp* mRNA for 12 h. **H** MATR3 (red) and GDF9 (green) staining in GO which had knocked down MATR3 protein before injected *Matr3-egfp*, *Δrrm2-egfp*, *Δpnls4-egfp*. n ≥36. **I** Quantification of the mean fluorescence intensity of GDF9 (**H**) in oocytes. Scale bar: 40 μm. Data are represented as mean±SD. \*\*\**P* < 0.001, n.s., not significant.

We injected low-concentration of *Matr3-Egfp* mRNA into either GOs or FGOs and performed live-cell fluorescence imaging. The results showed that in GOs and NSN-type FGOs, MATR3 was localized in the nucleus, while in SN-type oocytes, MATR3 presented as aggregated punctate structures and diffuses into the cytoplasm as meiosis resumes (Fig.5F). In addition, we injected the truncated form into germinal vesicle (GV) oocytes to further detect the localization of MATR3 (Fig.S8C). The CO - IP results indicated that MATR3 interacts with KDM3B through the 2nd RNA recognition motif (RRM2) and the 4th nuclear localization signal (pNLS4) (Fig.5E). Furthermore, we injected *in vitro*-transcribed mRNAs (*egfp*, *Matr3-egfp*, *Δrrm2-egfp*, *Δpnls4-egfp*) into GOs and carried out living-cell fluorescence imaging. The results showed that the functional truncation mutants of the RRM2 motif and the pNLS4 motif both change the spatiotemporal specific localization of MATR3 (Fig.5G).

To determine the domain that affects the physiological function of MATR3 in oocytes, we firstly knocked down the endogenous level of MATR3 in GOs and then injected *Matr3-egfp*, Δ*rrm2-egfp*, and *Δpnls4-egfp* mRNAs respectively so as to detect the protein level of GDF9. The results showed that after deleting RRM2, there was no significant change in the GDF9 protein level, while after deleting the pNLS4 domain, the GDF9 level decreased significantly (Fig.5H, I). Together, these results suggested that MATR3 is localized in the nucleus of GOs through the pNLS4 domain, thereby recruiting KDM3B and promoting the expression of GDF9.

### 6. MATR3 promotes the expression of *Rdx* to facilitate GDF9 secretion

As a key OSF of oocyte paracrine, both the transcription and secretion of GDF9 is regulated by MATR3. Notably, our immunofluorescence results showed that the structure of Oo-Mvi in cKO mice was damaged (Fig.6A B), which may impair the communication of oocyte and the surrounding cumulus cells.

**Fig 6.**
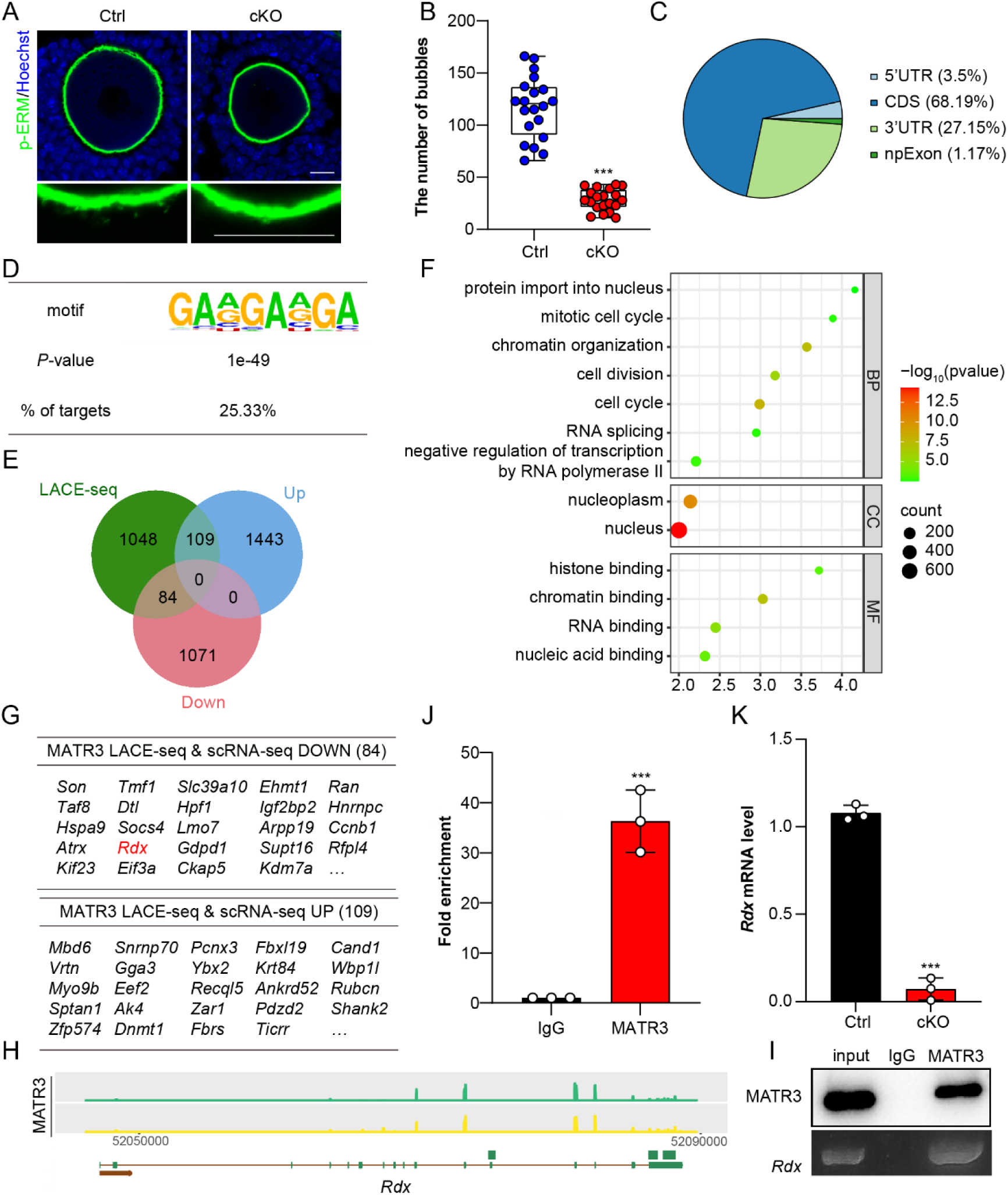
MATR3 is required for the OO-Mvi in mouse oocyte by regulating *Rdx* mRNA level. **A** p-ERM staining (green) showing the OO-Mvi in AFs from Ctrl and cKO. **B** Quantification of the number of Oo-Mvi vesicles (n = 20). **C** Pie chart showing the proportion of the reads values of 10-12-day-mice MATR3 LACE-seq. **D** Table showing the base sequences to which MATR3 mainly binds. **E and G** Venn diagrams (**E**) and table (**G**) showing overlapping genes binding with MATR3, upregulated genes and downregulated genes in scRNA-seq. **F** Key GO enrichment of all DEGs in LACE-seq. **H** IGV snapshot of MATR3 peaks distribution in 10-12-day-mice ovaries. **I and G** RIP showing the interaction between MATR3 and *Rdx* mRNA in 10-12-day-mice ovaries. Western blotting showing the reliability of MATR3 antibody (**I**). Enrichment degrees of MATR3 on the *Rdx* mRNA respectively (**J**). IgG served as the negative control and Input (2%) served as the positive control. n = 3. **K** RT-qPCR results showing *Rdx* mRNA level in PD14 oocytes. Scale bar: 20 μm. Data are represented as mean±SD. \*\*\**P* < 0.001.

To test if MATR3 in GO affects oocyte-somatic cell communication by binding to the DNA sequences pivotal for oocyte growth, the low-input affinity cleavage enrichment sequencing (LACE-seq) assay was performed by collecting growing oocytes from 10 to 12-day-old mice. The sequencing results showed that MATR3 directly binds to the coding sequence (CDS) region and 3’ untranslated region (3’UTR) of mRNAs (Fig.6C). Motif analysis indicated that MATR3 tends to bind to regions rich in GA (Fig.6D). The results of GO enrichment analysis showed that the genes bound by MATR3 were significantly enriched in biological processes such as nucleocytoplasmic protein shuttling, chromatin assembly, and RNA splicing (Fig.6F).

Later, a joint analysis of LACE-seq and scRNA-seq was performed to further confirm our assumption that MATR3 is very important for oocyte-cumulus cells communication. The results showed that among the genes directly bound by MATR3, 84 genes such as *Son*, *Ehmt1*, *Ran*, *Igf2bp2*, *Ccnb1*, and *Rdx* simultaneously showed a significant decrease while 109 genes simultaneously showed a significant increase in the oocytes of the knockdown group (Fig.6E, G). Of which, RDX is a specific Oo-Mvi-related protein in oocytes that contributes to the formation of Oo-Mvi in oocyte development. The sequencing data were visualized by IGV. The results showed that MATR3 directly binds to *Rdx* mRNA (Fig.6H). Fifty of the ovaries of 10 to 12-day-old-mice were collected for RNA immunoprecipitation (RIP). Afterward, RIP was used to verify the direct binding of MATR3 to *Rdx* mRNA in the ovaries. Consistent with the sequencing results, MATR3 directly binds to *Rdx* mRNA (Fig.6I, J). Furthermore, the *Rdx* mRNA level in the GOs of the cKO group was significantly decreased (Fig.6K), respectively, as compared with the mice in the Ctrl group. In sum, the above results indicates that in GOs, MATR3 directly binds to *Rdx* mRNA and regulates its level, thereby participating in the GDF9 secretion through Oo-Mvi.

## Discussion

This study shows that the spatiotemporal-specific localization of MATR3 in GO is required for female fertility. MATR3 in GO not only affects the transcription of specific genes but influences the overall chromatin organizational conformation. Further, in the condition that Matr3 is deleted *in vivo* or is silenced *in vitro*, seldom of growing follicles could develop into antral follicles. Consequently, the efficiency of superovulation is markedly downregulated, indicating a poor response to gonadotropins in cKO mice. *In vitro*, GDF9 partially rescued the phenotype of antral follicle failure as well as oocyte maturation caused by si-*Matr3* implying that GDF9 is one of the downstream molecules of MATR3. Furthermore, reduced GDF9 and RDX productions induced by MATR3 missing impairs the mutual communication between GO and surrounding granulosa cells. In agree with the existed reports, this is one of typical examples to emphasize the importance of OSFs on granulosa cell proliferation as well as GO growth (Gilchrist et al. 2008). We further demonstrated that MATR3 may exert its action by either coordinating KDM3B directed histone methylation, chromosome structure loosing and improving gene transcription through directly binding to the promoters of targeted genes. Since the protein expression pattern of MATR3 in oocytes of human, porcine and mice are similar to each other, MATR3 is a conservative key regulator of GO growth and follicle development among mammals (Fig.7). Therefore, MATR3 is a key regulator of oocyte growth and follicular development.

**Fig 7.**
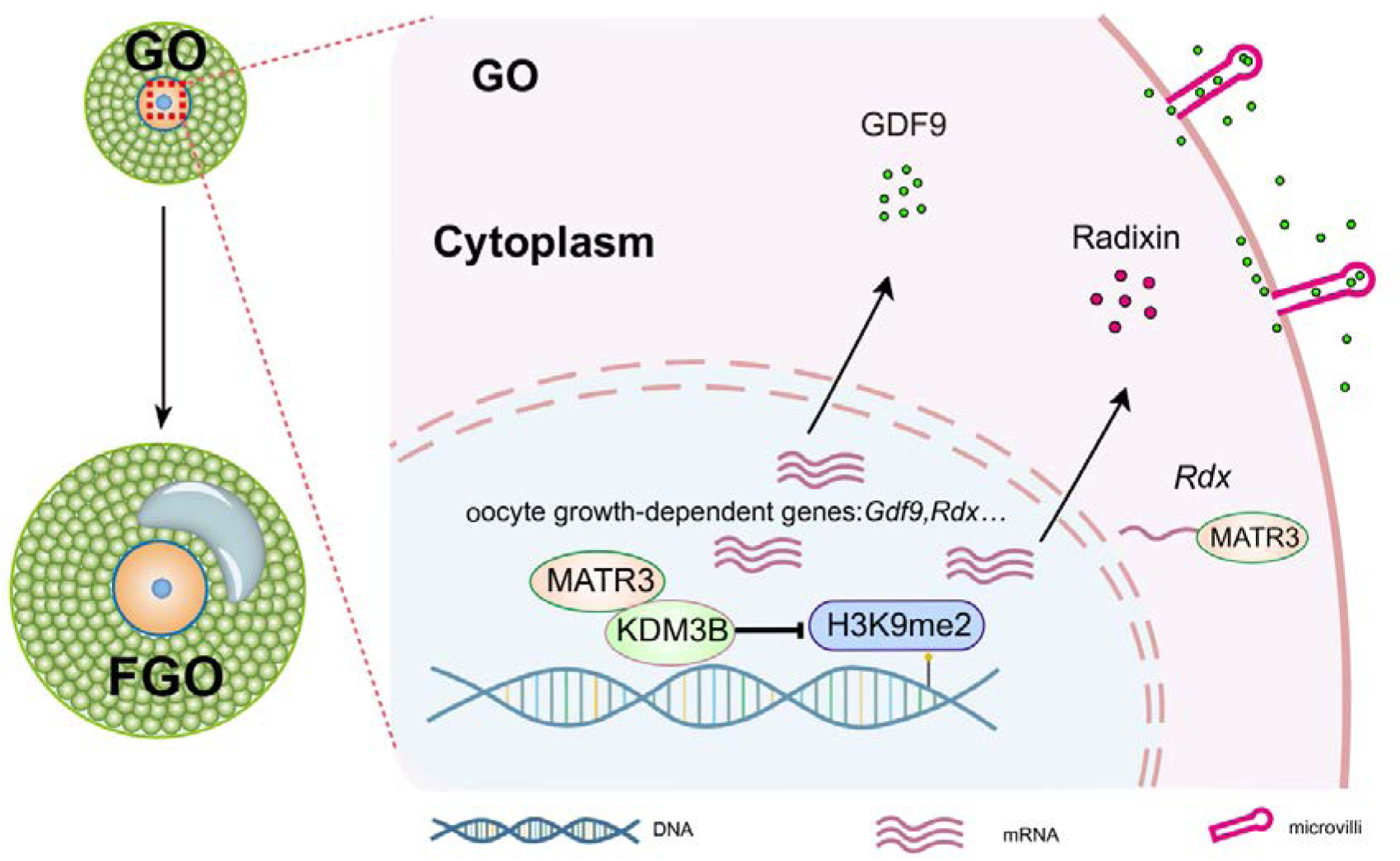
A proposed model: MATR3 regulates the synthesis and secretion of OSFs, like GDF9, thereby ensuring GO growth. Under physiological conditions, MATR3 in the growing oocytes promotes the transcription of a large number of genes required for cell growth. When promoting the transcription of the *Gdf9* gene, MATR3 localizes in the nucleus through the pNLS4 domain to recruit KDM3B, thereby promoting the high-level expression of *Gdf9*. In ensuring the secretion of GDF9, MATR3 directly binds to *Rdx* mRNA to promote the growth of microvilli, which in turn lays the foundation for ensuring the secretion of factors such as GDF9 and establishing the physical connection between oocytes and somatic cells.

During follicle growth, the high transcriptional activity in GO is a prominent physiological feature of oocytes. The uniqueness of GOs lies in their high transcriptional activity, if compared with oocytes at other developmental stages. When primordial follicles are activated, oocytes in the follicles are transformed from dormant state to the growing state. At this time, a large number of mRNAs start to be transcribed within the oocytes. As GO progress to FGO, transcriptional silencing occurs, which persists until the 2-cells stage. This indicates that the synthesis and accumulation of mRNAs for subsequent oocyte maturation, fertilization, and early embryo development mainly occur in the GO. Consequently, the large amounts of maternal mRNAs, proteins, and metabolites accumulated in the cytoplasm of FGO are important reasons for the gradual increase in oocyte volume, and are also essential for the oocytes to acquire the abilities of meiosis, fertilization, and embryo development (Su et al. 2021). Therefore, the high transcriptional level of mRNAs during the growth process of GOs is of particular importance.

MATR3 could also be one of candidate causative RBPs contributing to OMA in females. Recently, mutations of some RBPs pivotal for maternal mRNA homeostasis have been linked to the consequence of OMA, including PATL2 (Hu et al. 2024), PABPC1L (Wang et al. 2023) and ZFP36L2 (Wan et al. 2024). Here, the localization of MATR3 changes in oocytes of OMA patient as well. Specifically, MATR3 is highly expressed in the nucleus of NSN oocytes and exits the nucleus when the nuclear type transforms to SN in healthy people. However, in an OMA oocyte that remained at GV stage after ICSI, the nuclear localization of MATR3 disappeared. Moreover, the diameter of this oocyte was smaller than that of normal FGO. Furthermore, we noticed that deletion of MATR3 in mouse GOs resulted in significant chromosome configuration disorders by that the percentage of SN versus NSN FGOs was reversed in cKO mice. These findings imply that MATR3 is pivotal for chromosome configuration. In agree with existed studies explaining that the SN chromatin configuration correlates with transcriptional silence and full oocyte developmental competence (Bouniol-Baly et al. 1999; Zuccotti et al. 2005), this study also found that multiple oocyte-specific genes and genes involved in oocyte/granulosa cell communication are direct targets of MATR3, including *Ehmt1* (Demond et al. 2023), *Ran* (Dehapiot and Halet 2013), *Igf2bp2* (Li et al. 2021), *Ccnb1* (Wang et al. 2024b) (Fig.S9), *Rdx* (Zhang et al. 2021), *Zar1* (Rong et al. 2019) and *Dnmt1* (Shelby et al. 2023). The transcription of these genes changed in the condition of MATR3 loss. Therefore, MATR3 is pivotal for chromosome configuration and global gene transcription in GO, which is meaningful for the growth of oocyte. Despite these findings, however, it is too early to treat MATR3 as one of biomarkers to OMA since we could not collect more oocytes from OMA to confirm the conclusion so far. More clinic studies are needed to give confirmed relationship between MATR3 and OMA. Possibly, analysis of any mutations or irregular expression levels of MATR3 in immature oocytes of OMA patients may be pivotal to confirm the causal relationship.

This study provided additional evidences to highlight the crucial roles of OSFs in supporting follicular development at different stages (Dong et al. 1996). Studies including ours have shown that GDF9 is indispensable for granulosa cells proliferation during the transition from primary follicles to secondary follicles (Dong et al. 1996; Ackert et al. 2001; Gao et al. 2024). In the differentiated cumulus cells of antral follicles, GDF9 inhibits the synthesis of estrogen and the expression of LH receptors, thereby maintaining the differentiated state of cumulus cells. Meanwhile, it promotes the expression of NPR2 in cumulus cells, upregulates the levels of cGMP and cAMP in oocyte, and contributes to the oocyte meiotic arrest (Hinckley et al. 2005; Norris et al. 2009; Zhang et al. 2010). *In vitro*, GDF9 promotes the synthesis of new transzonal projections in the cumulus-oocyte complex (El-Hayek et al. 2018). Besides the powerful roles in regulating multiple gene expression responsible for oocyte development and oocyte/granulosa cell mutual communication, this study specifically emphasized the importance of MATR3 controlled production and secretion of GDF9 on antral follicle formation as well as oocyte maturation with the assistance of RDX for establishing profound Oo-Mvi. This is one of typical examples to emphasize the importance of OSFs on granulosa cell proliferation, GO growth as well as antral follicle formation, which are all pivotal for supporting high qualified oocyte development. This study highlights the importance of a sustaining feedback regulation between the oocyte and the surrounding somatic cells on GO growth and follicle development.

Previous studies reported that GOs could be divided into three stages based on the diameter, namely GO1 (30 - 45 μm), GO2 (50 - 55 μm), and GO3 (60 - 65 μm) (Gu et al. 2019). According to our results, there is no significant difference in the abundance of transcripts and chromatin accessibility among the three states of GO, but the DNA methylation levels vary significantly. After knocking down *Matr3* in GO2, the levels of H3K4me3 and H3K27me3 decreased significantly, which is contrary to the increasing trend of H3K4me3 and H3K27me3 levels with oocyte growth depicted in the DNA methylation map. The results proves that MATR3 is a positive regulatory on GO growth. During the GO growth, the levels of classical histone methylation modifications H3K4me3 and H3K9me3 increase and reach their peaks in FGO (Kageyama et al. 2007). Here, the levels of H3K9me1 and H3K9me3 linked to gene transcriptional repression were unchanged while H3K27me3 and H3K4me3 decreased significantly, indicating that classical histone methylations do not play dominant roles in MATR3-mediated transcription in GO. Instead, the level of H3K9me2, which is related to gene transcriptional repression, increased significantly. Consistently, the level of KDM3B who directly regulates H3K9me2 decreased significantly (Kim et al. 2012). So, H3K9me2 is crucial for MATR3-mediated high transcriptional levels in GO. Specifically, the pNLS4 domain of MATR3 recruits KDM3B, and regulate the transcriptional synthesis of GDF9 by removing the inhibitory H3K9me2 mark. Besides its coorperation with KDM3B to regulate gene transcription ability, MATR3 may directly bind to the promoters of target genes like *Rdx* to regulates transcription.

Overall, this study identifies MATR3 as a pivotal RBP essential for oocyte growth and female fertility, with conserved roles across multiple species including human, porcine, and mouse. We demonstrate that MATR3 operates through a dual molecular mechanism: it epigenetically promotes the transcriptional synthesis of *Gdf9* by recruiting the H3K9me2 demethylase KDM3B, and post-transcriptionally facilitates GDF9 secretion via binding and stabilizing *Radixin* mRNA. Disruption of this coordinated regulatory network impairs granulosa cell communication and arrests follicular development at the secondary stage, ultimately leading to infertility. Given its functional conservation and central role in oocyte maturation, MATR3 represents a promising diagnostic marker and therapeutic target for clinical OMA.

## Materials and methods

### Animals

*Gdf9*-Cre mice were generated by Prof. Fengchao Wang (Transgenic Animal Center, National Institute of Biological Sciences, Beijing, China). *Matr3* ^LoxP/LoxP^ mice were generated by Prof. Fengchao Wang using CRISPR-Cas9 technology. The sgRNAs were prepared using MEGAshortscript T7 Transcription kit (Ambion) according to the manufacturer’s instructions. Cas9 protein, sgRNAs and donor templates were injected into C57BL/6J fertilized eggs. Injected zygotes were transferred into pseudo-pregnant CD1 female mice. Sequence of gRNA and primers used for genotyping were listed in Table S1.

### Mouse fertility and ovulation assay

For the mice fertility test, a 2-month-old cKO female and its littermate control (Ctrl) female were housed with a 2-month-old C57BL/6J male with normal fertility. Each male mouse was housed with one cKO and one Ctrl female mouse simultaneously. Mating cages were monitored daily. The number of pups (both alive and dead) was counted on the first day of delivery. The mating process lasted for 6 months.

For superovulation, cKO and Ctrl mice at post parturition (PD) 23 days were intraperitoneally injected with 5 IU PMSG (Ningbo Sansheng Biological Technology, Cat#110251283), followed by 5 IU hCG (Ningbo Sansheng Biological Technology, Cat#110041282) 46 h later. After an additional 13 h, oocytes of each mouse were collected from oviducts, and the number of oocytes was counted after digesting with 0.3% hyaluronidase (Merck, Cat#MR-051-F).

### Follicle counting

Fresh ovarian samples were fixed in 4% paraformaldehyde (Santa Cruz, Cat#30525-89-4) overnight, embedded in paraplast (Leica, Cat#39601095), and sectioned serially at 8 μm. Tissue sections were stained with hematoxylin (Solarbio, Cat#G4070) to count the number of follicles. Sections were examined and photographed using VENTANA DP200 (Roche).

We classified the follicle stage by its morphological characteristics. A PrF contained an oocyte with a diameter less than 20 μm and one-layer flattened pre-granulosa cells (GCs); A PF contained a larger oocyte and one-layer cubical GCs; A SF had multilayer GCs; An AF had antral cavity.

The number of every follicle stage was summarized by counting all sequential sections. In both cases, only follicles containing clearly visible oocyte nucleus in each individual section were counted to avoid repetitive counting.

### Immunostaining and biological assays

The ovarian sections were deparaffinized, rehydrated, and subjected to high-temperature (95–98 °C) antigen retrieval for 16 min with 0.01% sodium citrate buffer (pH 6.0). Immunohistochemistry assay was performed using Histostain™-SP Kits (ZSGB-BIO, Cat#PV-9001) and DAB peroxidase substrate kits (ZSGB-BIO, Cat#ZLI-9017) according to the manufacturer’s protocols. Nuclei were stained with hematoxylin (Solarbio, Cat#G4070). Primary antibodies and dilution rates were as follows: rabbit anti-MATR3 (1:400, Abcam, Cat#ab151714) Sections were examined and photographed using VENTANA DP200 (Roche).

For immunofluorescence assay, ovarian paraffin sections were deparaffinized, rehydrated, and subjected to high-pressure antigen repair with 0.01% sodium citrate buffer (pH=6.0) for 16 min. The sections were then rinsed thoroughly with phosphate-buffered saline (PBS) for 10 min and blocked with 10% normal donkey serum (Yesean, Cat#36116ES10) in PBS for 1 h at room temperature and incubated with primary antibodies (diluted with PBS) for 16 h at 4°C. Primary antibodies and dilution rates are as follows: mouse anti-DDX4 (1:400, Abcam, Cat#ab27591); goat anti-FOXL2 (1:400, Novus, Cat#NB100-1277); rabbit anti-p-ERM (1:300, Cell Signaling Technology, Cat#3726T); rabbit anti-SMAD3 (1:200, Cell Signaling Technology, Cat#9523); rabbit anti-Ki67 (1:400, Cell Signaling Technology, Cat#D385). Next, ovarian sections were rinsed thoroughly with PBS for 1 h and incubated with Alexa Fluor 488- or 555- conjugated secondary antibody (1:200, Yesean, Cat#33106ES60) for 1 h at 37°C. Subsequently, the sections were again rinsed thoroughly with PBS, stained with Hoechst33342 (1:100, Sigma, Cat#14533) for 1 min, and sealed in anti-fade fluorescence mounting medium (Applygen, Cat#C1210) with microscope cover glass (Citoglas, Cat#10212450C). Sections were examined and photographed using a Nikon A1 Confocal microscope.

For Ki67-positive GC analysis, the percentage was quantified as the number of GCs with positive signal divided by the total number of GCs per maximum follicular cross-section in the SFs of Ctrl and cKO mice. Sections were examined and photographed using a Nikon A1 Confocal microscope.

### Early growing follicle isolation and culture *in vitro*

10-12 old days ICR mice were obtained from Beijing Vital River Laboratory Animal Technology Co., Ltd. and housed at China Agricultural University. To collect the follicles, ovaries were dissected, discarded the interstitial tissues and incubated in modified Leibovitz’s L-15 medium (Gibco 11415064) added with 1×penicillin-streptomycin (Gibco 15070063, 100×). The pre-antral follicles with indistinguishable antral structure based on relative size of 150 - 200 μm were confirmed using stereomicroscope. Under the stereomicroscope, the ovaries were cut into small pieces using a 1 mL insulin syringe needle with the “cross-shaped division method”. Early growing follicles with a diameter of approximately 150 - 200 μm and 5 - 6 layers of granulosa cells were selected under the microscope, and then isolated from the ovarian tissue as intact individual follicles using the insulin needle. Early growing follicles for each experiment were collected from 2 - 3 mice. At least 20 - 30 follicles were collected for each group.

Isolated early growing follicles were cultured on 0.4 µm pore size inserts (Millipore, Cat#PICM0RG50) in 6-well culture plates (NEST, Cat#703002) for 5 days. Basal culture medium comprised 1.6 mL MEM-α medium, 1×insulin-transferrin-sodium selenite media supplement (Sigma I3146, 100×), 25 mM NaHCO3, 5% fetal bovine serum (Gibco 16000-044), 1×penicillin-streptomycin and 10 ng/mL follicle stimulating hormone (FSH) purchased from National Hormone and Peptide Program. As a control, follicles were microinjected *Matr3* siRNA (20 μM, Sigma-Aldrich, Cat#EMU014021) or negative control siRNA (20 μM, GenePharma). Approximately half of the medium in each well was replaced with fresh medium every other day. Follicles were maintained at 37°C under 5% CO2 and 95% air.

### Oocyte isolation and culture *in vitro*

For GOs, the ovaries consisted with pre-antral follicles of 10 - 12 days old mice were dissected in modified M2 medium (Millipore, MR-015-D). The ovaries were digested carefully in M2 medium containing 0.25 ng/mL collagenase type IV (Sigma-Aldrich, C5138), and the isolated GOs were cultured in M2. Upon the overnight culture in the M2, GOs containing an intact GV and indistinguishable zona pellucida were used for further assays.

For FGOs, mice at PD23 received intraperitoneal injections of 5 IU PMSG (Ningbo Sansheng Biological Technology, Cat#110251283). After 46 h, the FGOs were released into the M2 medium. The germinal vesicle (GV) oocytes were then cultured in drops of M16 medium (Millipore, MR-016) covered with mineral oil at 37◦C. Germinal vesicle breakdown (GVBD) and the extrusion of the first polar body (MII) were observed and recorded at 2 and 16 h, respectively.

### Plasmid construction

Full-length mouse *Matr3* cDNA was obtained from GO and cloned into the pcDNA3.1(+) vector using BamHI and XhoI endonuclease. To generate MATR3-EGFP, the EGFP open reading frame with a 14 amino acid N-terminal linker was amplified from pLV-EGFP-Cre by PCR using forward primer 5’-AAG GAA ACT GTG AGC AAG GGC GAG GAG C −3’ and reverse primer 5’- AGT GGA TCC GAG CTC GGT ACC CTA CTT GTA CAG CTC GTC CAT GCC −3’. To create pcDNA3.1(+)-MATR3-EGFP, the MATR3 open reading frame from pCDNA3.1(+)-MATR3 was amplified by PCR with forward primer 5’- GGG AGA CCC AAG CTG GCT AGC ATG TCC AAG TCA TTC CAG CAG TC-3’ and reverse primer 5’-CCC TTG CTC ACA GTT TCC TTC TTC TGC CTC CG-3‘, digested with NheI and KpnI, and inserted into the corresponding sites in pcDNA3.1(+). All constructs were confirmed by sequencing prior to transfection in HEK293T cells (Abcam Cat# ab282205, RRID: CVCL_0063). Domain deletion mutants were described in previous research.

### *In vitro* transcription of mRNAs

Recombinant pcDNA3.1(+) plasmid containing the CDS of desired fragments were digested with XbaI. The DNA purification procedure was performed using TIANgel Midi Purification (TIANGEN BIOTECH, DP209) kit. Purified linearized plasmid templates were dissolved in RNase-free water and transcribed using the mMessage mMACHINE T7 ULTRA (ThermoFisher Scientific, AM1345) kit to generate the capped mRNA with poly(A) tails. For removing unincorporated nucleotides and most proteins, the mRNA synthesis reactions were stopped and precipitated with 50 μL lithium chloride precipitation solution at −20 ℃ overnight. On the following day, the mRNAs pellet was centrifuged to remove the unincorporated nucleotides and LiCl and resuspended in the RNase-free water. The final mRNA concentration was determined via Nanodrop (ThermoFisher Scientific). All the kits were used per manufacturer’s respective protocols.

### Oocyte immunofluorescence and confocal microscopy

The oocytes were fixed with 4% paraformaldehyde in PBS for 20 minutes at room temperature, followed by membrane permeabilized treatment with 0.5% Triton X-100 in PBS for 20 minutes at room temperature. After permeabilization, the oocytes were transferred into blocking buffer supplemented with 0.01% Triton X-100, 0.1% Tween 20 and 1% BSA for 1 hour at room temperature and the oocytes were then incubated with primary antibodies at 4 ℃ overnight. Primary antibodies and dilution rates are as follows: rabbit anti-MATR3 (1:400, Abcam, Cat#ab151714); rabbit anti-KDM3B (1:50, Abcam, Cat#ab70797); rabbit anti-p-ERM (1:300, Cell Signaling Technology, Cat#3726T); mouse anti-α-Tubulin-488 (1:100, Abcam, Cat#ab195887); mouse anti-H3K9me2 (1:200, Abcam, Cat#ab1220); rabbit anti-H3K27me3 (1:100, Active Motif, Cat#39155); rabbit anti- H3K9me1 (1:100, Abcam, Cat#ab176880); mouse anti- H3K9me3 (1:200, Active Motif, Cat#61013); rabbit anti- H3K4me3 (1:300, Abcam, Cat#ab8580); goat anti-GDF9 (1:100, Biotechne, Cat#AF739). Following by removal of the primary antibodies, the oocytes were washed three times in PBS washing buffer with 0.01% Triton X-100 and 0.1% Tween 20. For the secondary antibodies and staining, Alexa Fluor Plus 488 donkey anti-goat IgG (H+L) secondary antibody (Invitrogen, A32816) was used at a dilution of 1:100 for 2 hours at room temperature. After the secondary antibody incubation, oocytes were washed in washing buffer for three times and DNA was stained with Hoechst 33342 (1 μg/mL in blocking buffer) for 10 minutes at room temperature. Finally, the oocytes were dispersed and mounted on glass slides with DABCO-containing blocking buffer and analyzed by laser-scanning confocal microscopy.

### RT-qPCR

Total RNA was isolated from ovaries with TRIzol (Invitrogen, Cat#15596018). The quantity and quality of total RNA were determined using Nanodrop. 1 μg total RNA of each sample was used to reverse transcribe into cDNA according to manufacturer’s recommendation (Takara, Cat#RR047A).For oocyte, 50-100 oocytes were collected as one sample. Each sample was used to reverse transcribe into cDNA according to manufacturer’s recommendation (TRAN, Cat#AT301). RT-qPCR was performed in 96-well plates (Roche, Cat#04729692001) using FastStart Universal SYBR^®^Green Master (Roche, Cat#61396600) with LightCycler^®^ 96 Real-Time PCR System (Roche). Reaction parameters were as follows: 10 min at 95°C, followed by 45 cycles of 10 s at 95°C and 30 s at 60°C. Data were normalized to*β-Actin*. Primers are presented in Table S2.

### Western blotting

Total protein from ovaries was extracted with TRIzol (Invitrogen, Cat#15596018). For oocyte, 250 GOs or 200 FGOs were collected as one sample. Protein separated on 10% SDS-PAGE, and transferred to PVDF (polyvinylidene fluoride) membranes (Millipore, Cat# IPVH00). The membranes were blocked in 5% skim milk (Solarbio, Cat#D8340) for 1 h at room temperature and incubated with relevant primary antibodies (diluted with TBST (TBS plus 0.05% Tween-20)) overnight at 4°C. After rinsing with TBST, the membranes were incubated with the HRP-linked secondary antibody (1:4,000, ZSGB-BIO, Cat#ZB-2301/ZB-2305) for 1 h at room temperature and rinsed again with TBST. The membranes were visualized using the SuperSignal detection system (Thermo Fisher Scientific, Prod 34080). the image was quantified using Adobe Photoshop CS6. Primary antibodies and dilution rates were as follows: rabbit anti-MATR3 (1:1000, Abcam, Cat#ab151714); rabbit anti-KDM3B (1:500, Abcam, Cat#ab70797); rabbit anti-GAPDH (1:1000, Proteintech, Cat#10494); rabbit anti-RNAPII (1:10000, Abcam, Cat#ab5095); goat anti-FOXL2 (1:500, Novus, Cat#NB100-1277); rabbit anti-β-Actin (1:10000, Abmart, Cat#T40104); rabbit anti-GDF9 (1:500, Abcam, Cat#ab38544); rabbit anti-PCNA (1:500, Beyotime biotechnology, Cat#AF1363).

### scRNA-seq

GO of NC and si-*Matr3* were collected one by one, and RNA was extracted by commercial RNA extraction kit (Vazyme, Cat#N712). cDNA libraries were sequenced on the NovaSeq 6000 Illumina sequencing platform and analyzed by Novogene Co., Ltd. (Beijing, China). Differentially expressed genes (DEGs) were analyzed using the DESeq2 R package (1.20.0). Generally, genes with P values less than 0.05 and absolute fold-change larger than 2 were considered DEGs.

### LACE-seq

GOs were isolated from the ovaries of 10-12-day-old mice for LACE-Seq. Oocytes were washed three times with cold PBS, and immediately irradiated on ice with UV light at 400 mJ cm^−2^ for two times. The UV light treated oocytes were stored at −80 °C until 500 oocytes required for one experiment were accumulated. LACE-Seq was carried out as described previously, with the same experiment repeated independently twice.

### RNA immunoprecipitation (RIP)

Ovaries were isolated from the ovaries of 10-12-day-old mice for RIP. RIP assays were performed by using EZ-Magna RIP RNA-binding protein Immunoprecipitation Kit (Millipore; 17-701) following per manufacturer’s protocol. The first strand RIP cDNA libraries were synthesized with M-MLV synthesis kit (Invitrogen, C28025-032) by using random primers. Quantitative real time PCR assay was performed using FastStart Universal SYBR Green Master (ROX) (Roche, 04913914001) and primers related to for RT-PCR are shown in Table S1. The fold enrichment of *Rdx* in the anti-MATR3 antibody-precipitated ovaries was determined relative to IgG precipitated sample.

### CO-Immunoprecipitation

The antibodies and beads were combined overnight in a four-degree shaker. HEK293T cells (Abcam Cat# ab282205, RRID: CVCL_0063) were collected and lysate in RIPA lysate with PMSF. Then the supernatant was combined with beads which bind antibodies and kept overnight at 4℃. The beads were then eluted with the eluents in the Immunoprecipitation Kit (Invitrogen, 10006D). The precipitates were heated in the Sodium Dodecyl Sulfate-Polyacrylamide Gel Electrophoresis (SDS-PAGE) sample buffer and western blotting was performed.

### Statistical analysis

All experiments were repeated at least three times. Results were expressed as the mean ± S.D. Statistical analyses were conducted by GraphPadPrism9 software (GraphPad Software, La Jolla, CA, United States). The statistical significance of the differences between the groups was measured by two-sided ANOVA test. The statistical significances were defined as: *, *P* < 0.05; **, *P* < 0.01; ***, *P* < 0.001, n.s., non-significant.

## Supporting information

revised-Supplementary materials

## Acknowledgements

We appreciate the members of Wang’s and Xia’s laboratory for comments during preparation of the manuscript. This work was supported by the Beijing Natural Science Foundation (7252085), the National Key Research & Developmental Program of China (2024YFD1301001, 2023YFD1300501, 2022YFC2703803), the National Natural Science Foundation of China (32371167, 32071132, 32270904 and 32070839), the Innovative Project of State Key Laboratory of Animal Biotech Breeding (No. 2024SKLAB 1-1) and the 2115 Talent Development Program of China Agricultural University.

## Author contributions

YB, HZ and CW conceived the project. YB and CW designed the experiments. FW provided the *Matr3*^flox/flox^ and *Gdf9*-Cre mice. YB and TW managed conditional knockout mice. JM and RH provided clinical data. ZZ and LL supported the collection of oocytes. MG and SQ helped with the phenotypic analysis of knockout mice. QY, BL and WS helped with various experiments or analysis. GX, BZ, HZ, and CW supervised the project and related experiments. YB and CW wrote the manuscript with help from all authors.

## Data availability

All data generated or analyzed during this study are included in the article.

## Study approval

All experiments were endorsed by the General Hospital of Ningxia Medical University, No. KYLL-2021-758 and complied with the Declaration of Helsinki. Mice were housed in mouse facilities under 12/12-h light/dark cycles at 26°C and 40-70% humidity with access to chow and water *ad libitum*, according to the guidelines for the care and use of laboratory animals. All procedures were conducted in accordance with the guidelines of and approved by the Animal Research Committee of the China Agricultural University, No. AW71014202-3-1. All animal experiments involved ethical and humane treatment.

