## Supplementary material for "MATR3 is essential for oocyte growth and maturation quality through a dual molecular mechanism": revised-Supplementary materials

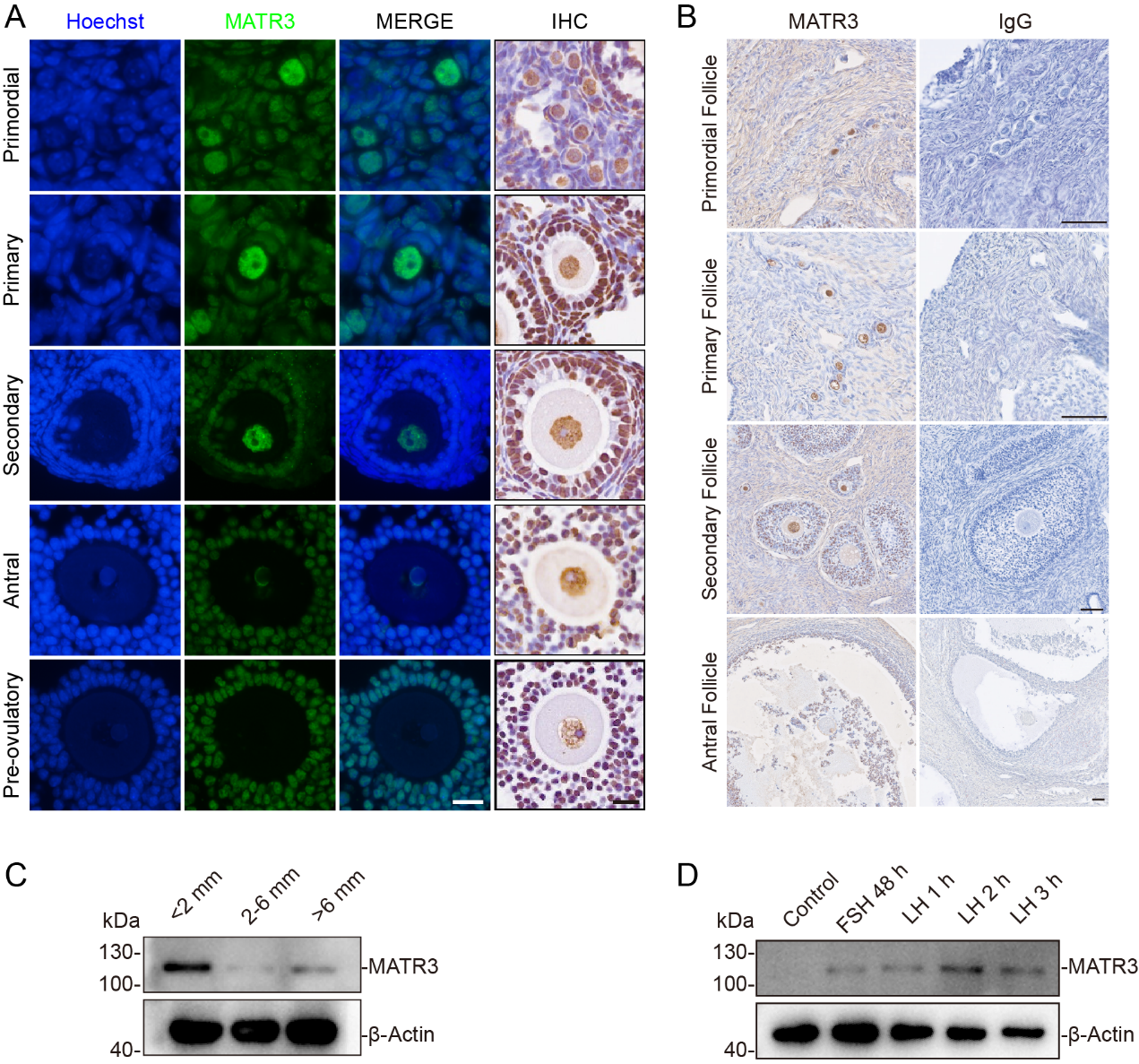
**Figure S1.** **The expression pattern of MATR3 in ovaries of mice and porcine.**

**A** Immunofluorescence results showing MATR3 location in mouse ovaries. Mouse ovaries were stained for MATR3 (green) at PD23.The nuclei were dyed with a Hoechst counterstain (blue). Scale bar: 25 μm. **B** Immunohistochemistry results showing MATR3 location in porcine ovaries. MATR3 was mainly localized to the nuclei of oocytes in either mouse or porcine. Scale bar: 50 μm. **C** Western blotting was used to detect the changes of MATR3 in GCs of porcine during the growth of follicles under physiological conditions. The results showed that MATR3 was highly expressed in the GCs of small secondary follicles, n = 3. **D** Western blotting was used to detect the changes of MATR3 in GCs of porcine with the duration of LH. The results showed that MATR3 increased with the duration of LH, n = 3.

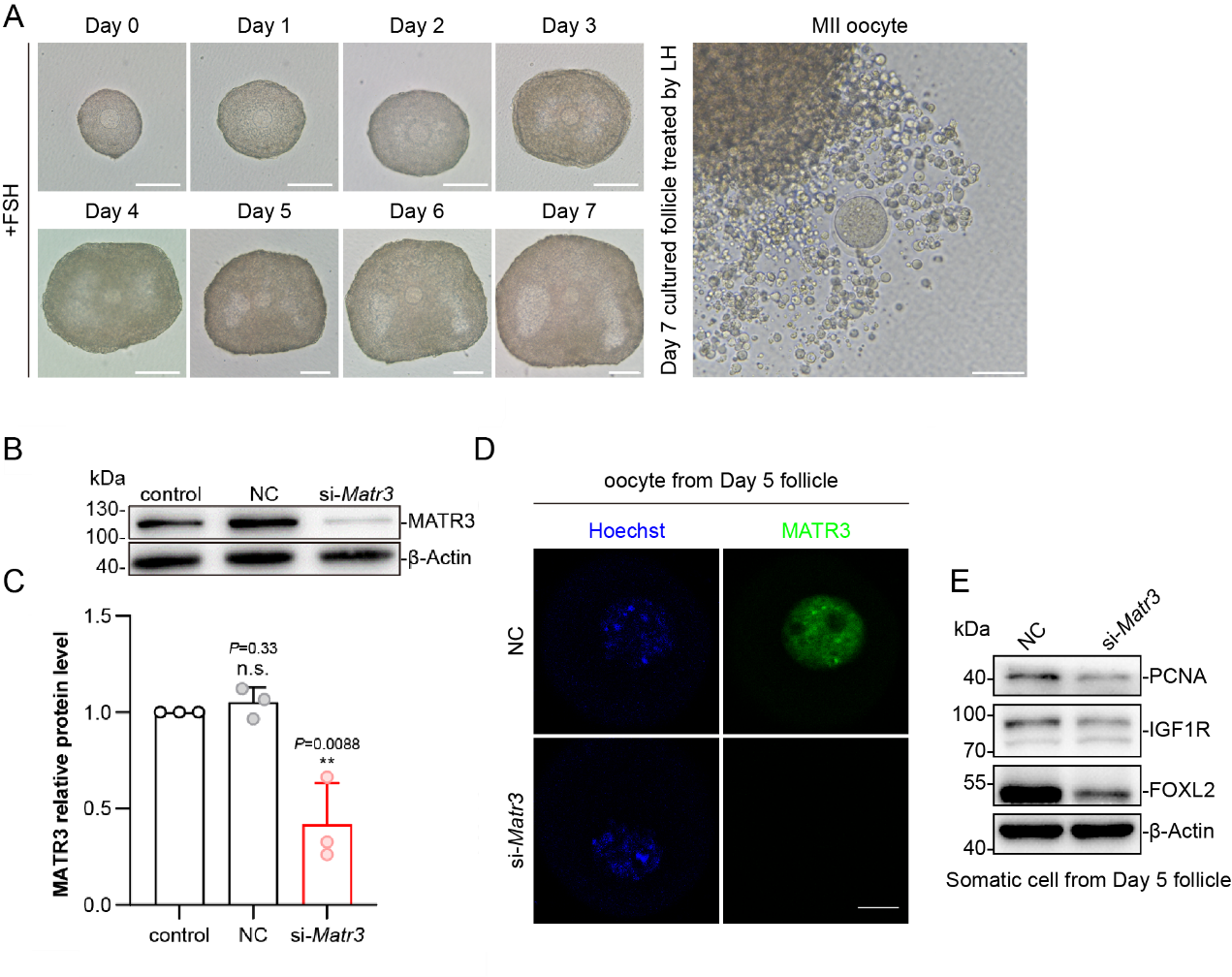

**Figure S2. Construction of *in vitro* knockdown model, knockdown efficiency, and phenotypes.**

**A** Using the method of *in vitro* addition of FSH, secondary follicles with a diameter of 150-200 μm were continuously cultured. The follicles could develop to the pre-ovulatory follicle stage, and oocytes with the ability to resume meiosis could be ovulated 16 hours after the addition of LH. Scale bar: 100 μm. **B** After the 3T3 cell line was transfected with NC and Matr3 siRNA respectively for 48 hours, Western blot was used to detect the protein level of MATR3 in the cells, with n = 3. β-Actin was used as an internal reference to normalize the protein loading level. **C** The statistical results of the gray scale scanning values of the MATR3 protein bands in (B). The statistical results are expressed as mean ± S.D. **D** The knockdown efficiency was detected by immunofluorescence technique. Green fluorescence: MATR3; Blue fluorescence: Hoechst (nucleus). Scale bar: 25 μm. **E** Isolate the somatic cells and oocytes of the follicles that have been continuously cultured for 5 days. Detect the changes in the levels of proteins related to the proliferation and differentiation of granulosa cells by Western blotting. n = 3.

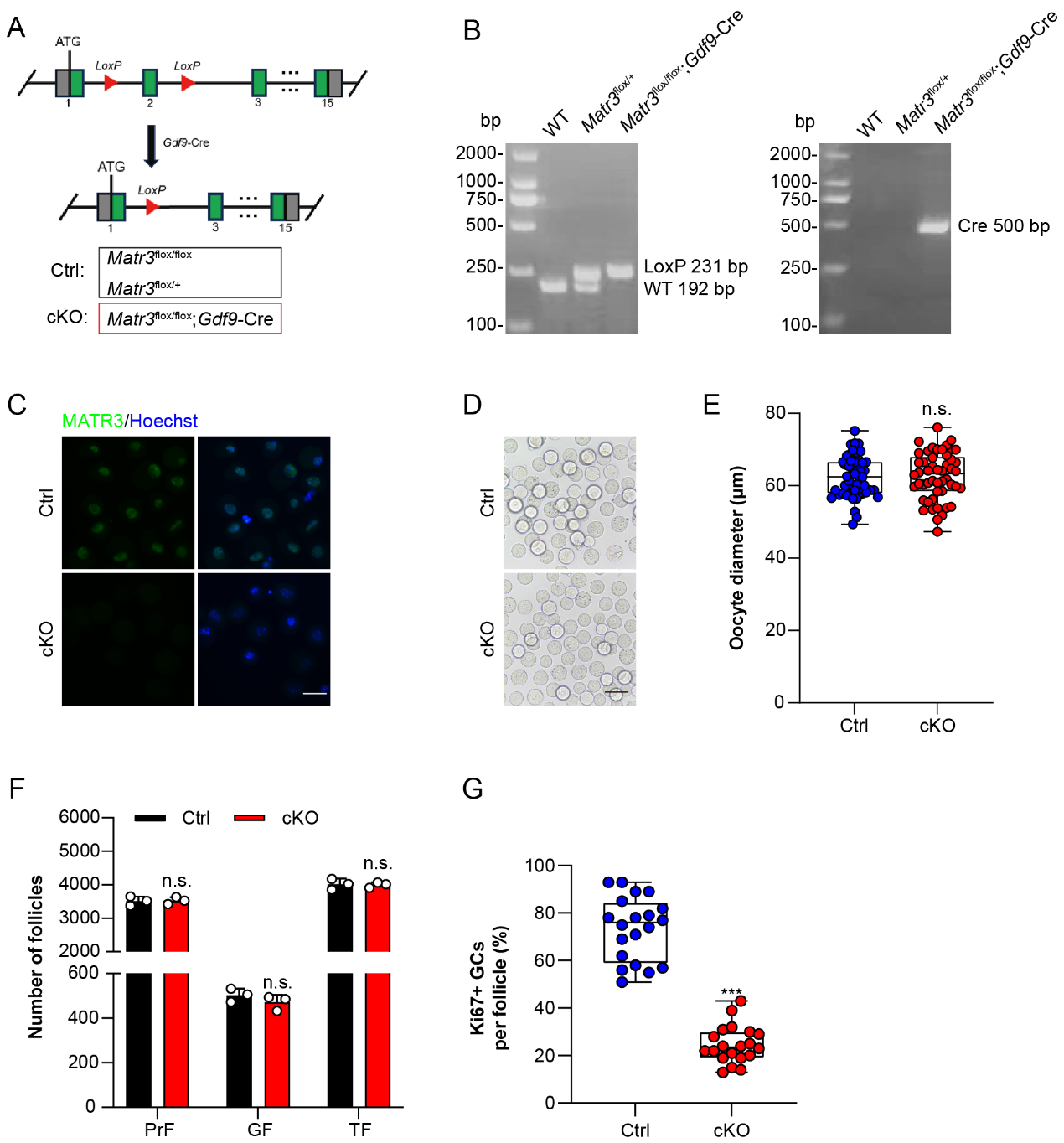

**Figure S3. Strategies for constructing *Matr3* oocyte-specific knockout mice and detection of related phenotypes.** **A** Construction strategy for *Matr3*-knockout model in oocyte via *Gdf9*-Cre. ATG: Transcription start site; Red arrow: LoxP insertion site; Numbers 1-15: Exon regions 1-15 of *Matr3*. **B** Using PCR to identify the genotypes of mice, with DL2000 as the DNA Marker. After the insertion of LoxP, the size of the band is 231 bp, the size of the wild-type (WT) band is 192 bp, and the size of the Cre is 500 bp. **C** Immunofluorescence results showing MATR3 location in oocytes from Ctrl and cKO mouse (PD14). Oocytes were stained for MATR3 (green). The nuclei were dyed with a Hoechst counterstain (blue). Scale bar: 50 μm. **D** Morphology of oocytes derived from PD14 mice. Scale bar: 50 μm. **E** Diameter of oocytes derived from PD14 mice. n = 50. **F** Number of follicles in PD7 ovaries. PrF: Primordial follicle; GF: Growing follicle; TF: Total follicle. **G** Quantification of the ratio of Ki‑67‑positive granulosa cells within individual follicles. n = 20. Data are represented as mean±SD. ****P* < 0.001, n.s., not significant.

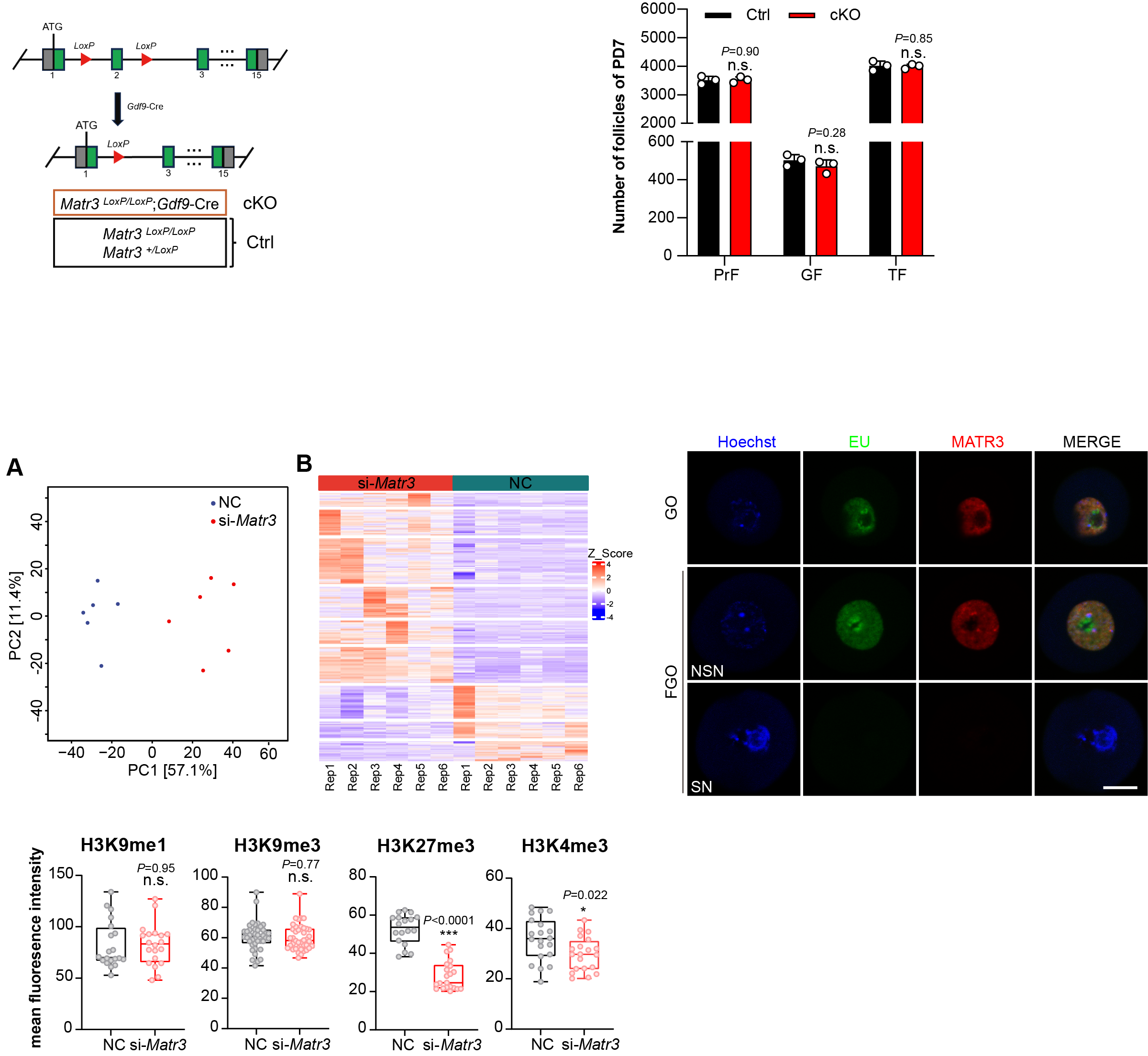

**Figure S4. The spatiotemporal-specific localization of MATR3 is adapted to the transcriptional activity of oocytes.**

Immunofluorescence results showing the relationship between the localization of MATR3 in oocytes at different growth states and the transcriptional activity of oocytes. Red: MATR3; Green: EU (newly synthesized mRNA); Blue: Hoechst (cell nucleus). Scale bar: 25 μm.

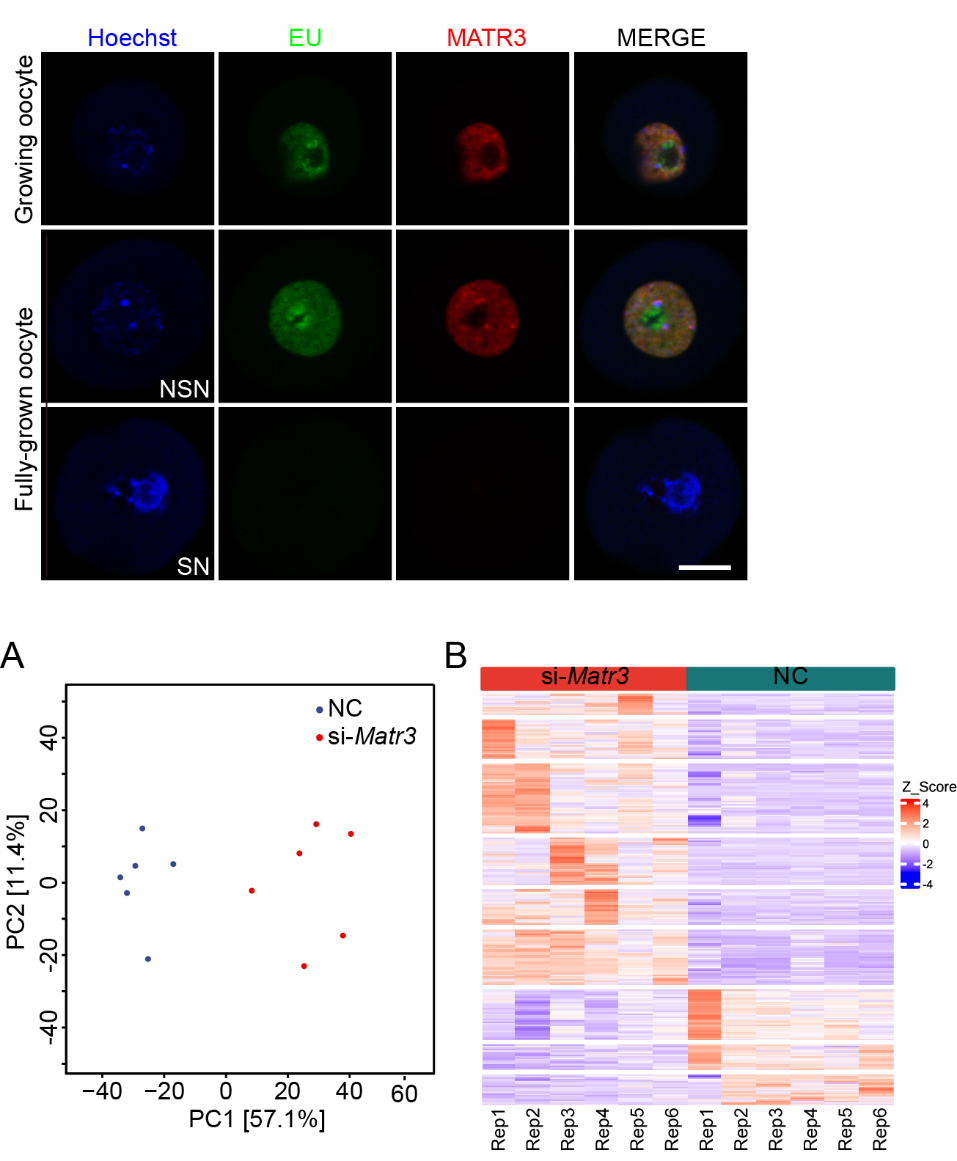

**Figure S5.** **Parallelism analysis of scRNA-seq samples.**

**A** PCA analysis shows that the GO in the NC group and the si-*Matr3* group form two subgroups. **B** Clustering heatmap analysis demonstrates the global changes in the expression levels of differentially expressed genes in the GOs of the NC group and the si-*Matr3* group based on the results of scRNA-seq data. The horizontal axis represents the types of sample replicates, and the vertical axis represents different genes.

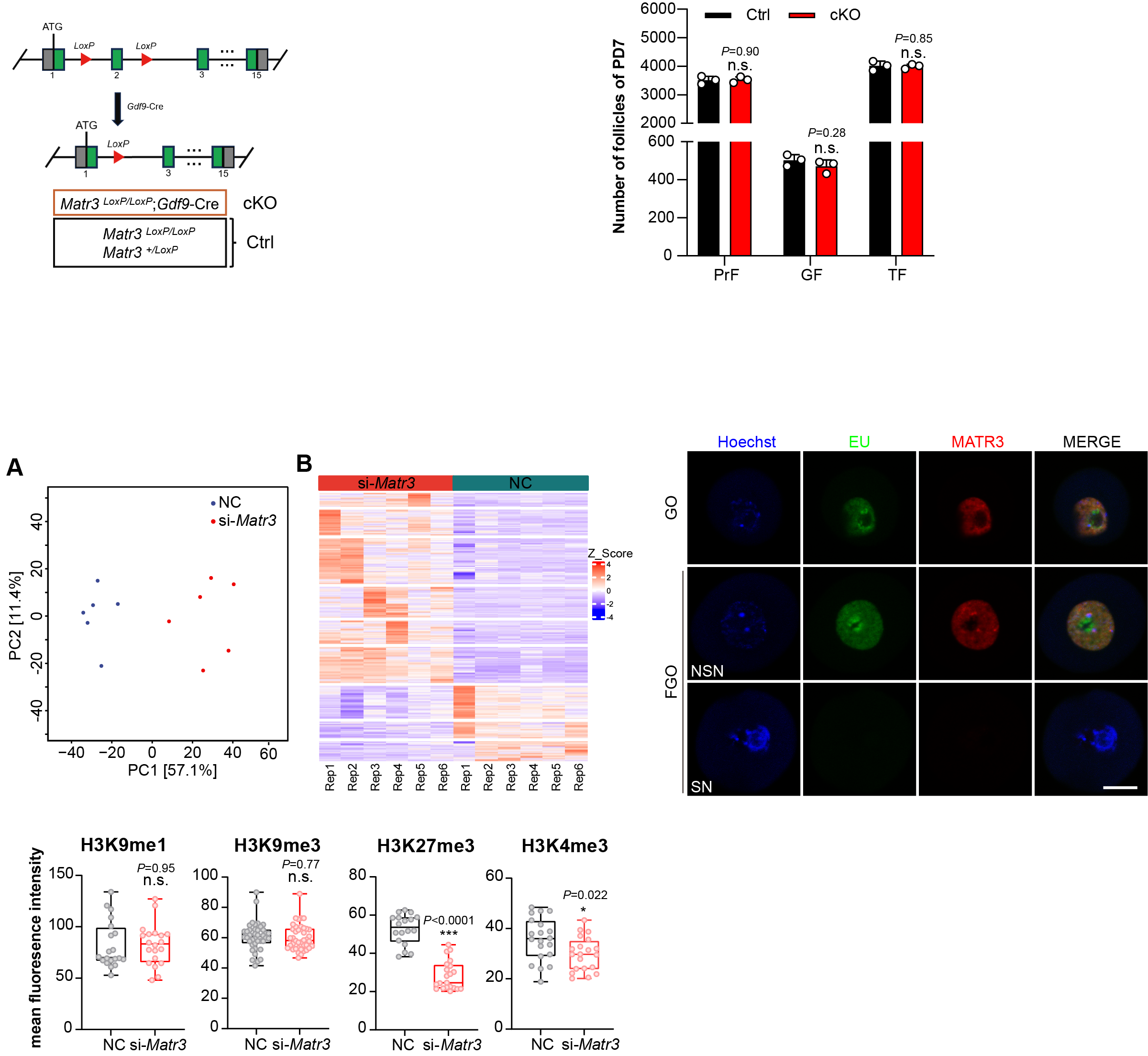
**Figure S6. The histone methylation level of oocytes after *Matr3* knockdown *in vitro*.**

The statistical results of the fluorescence intensity of histone methylation, with n (H3K9me1) = 20, n (H3K9me3) = 40, n (H3K27me3) = 18, and n (H3K4me3) = 20. The statistical results are presented as mean ± SD.

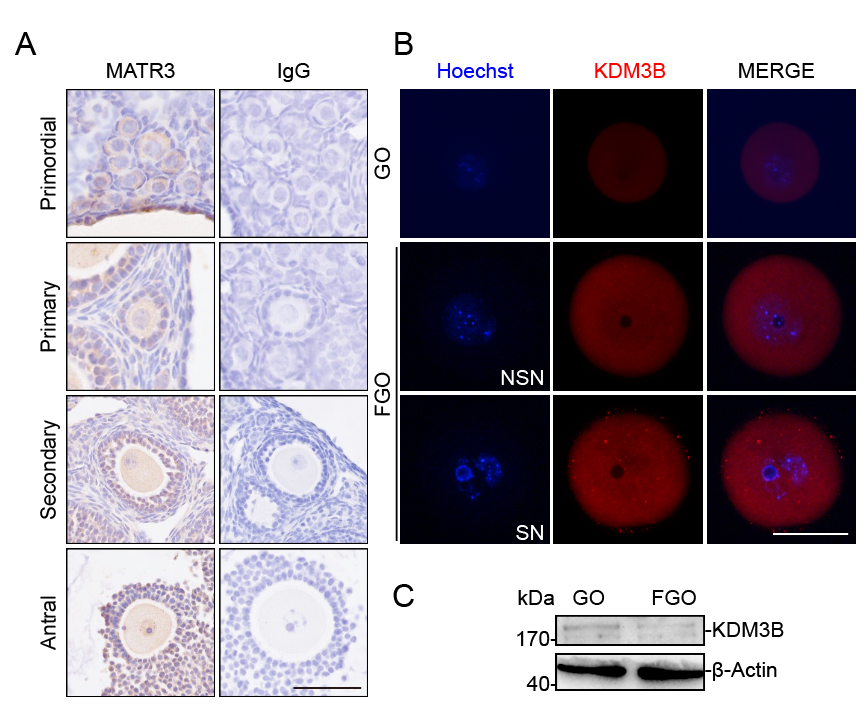

**Figure S7. The expression pattern of KDM3B in the ovaries of mice.**

**A** Immunohistochemistry results showing KDM3B location in mice ovaries. **B** Immunofluorescence results showing the location of KDM3B in GO and FGO. **C** Western blotting results showing KDM3B protein levels in GO and FGO. Scale bar: 50 μm.

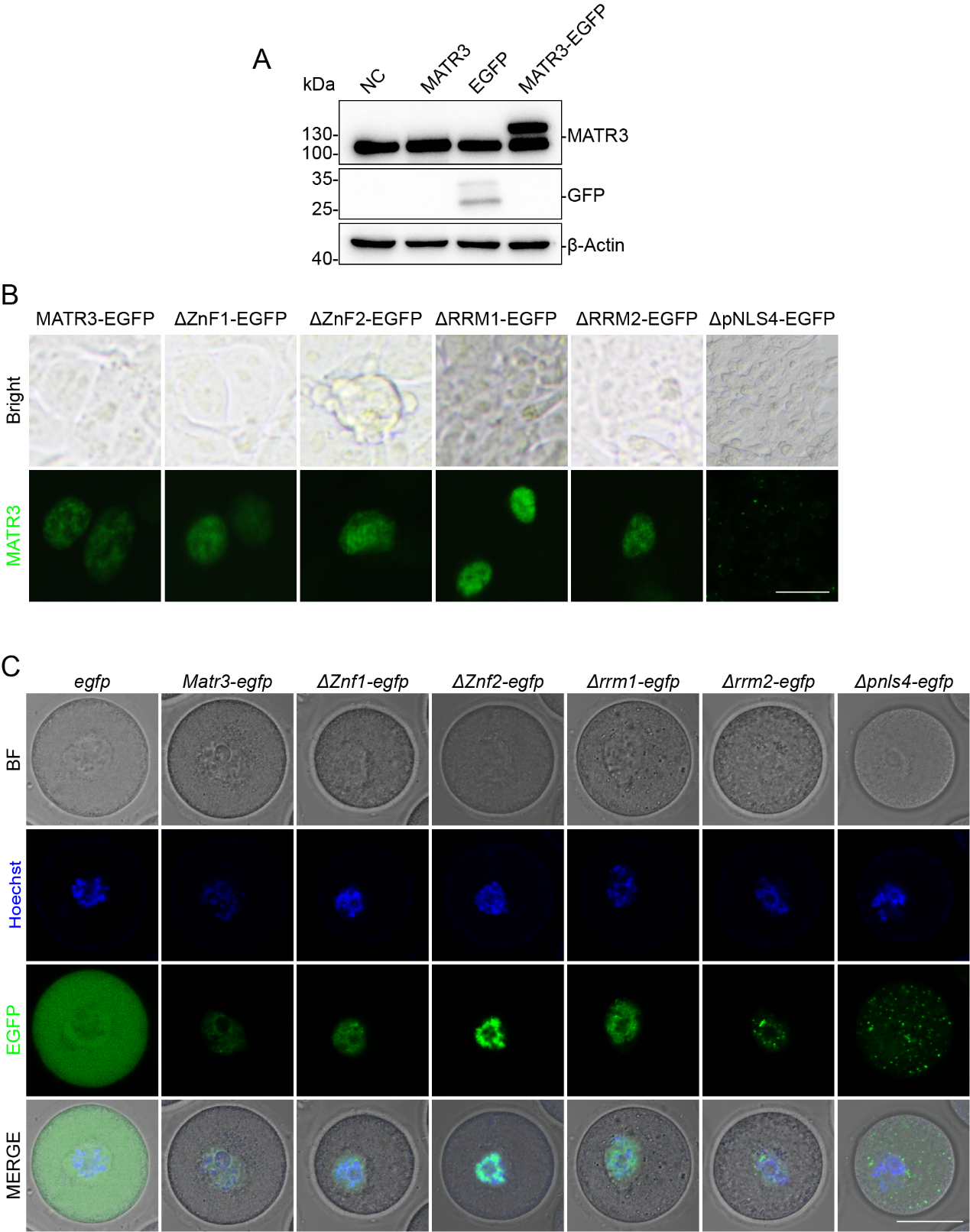

**Figure S8.** **Construction and localization detection of MATR3 truncated forms.**

**A** Western blotting was used to detect the protein expression levels of the 293-cell line after being transfected with NC, MATR3, EGFP, and MATR3-EGFP plasmids for 48 hours respectively. β-Actin is used as an internal reference to calibrate the protein loading level. **B** Immunofluorescence was used to detect the protein localization of 293 cell line after transfection with MATR3-EGFP, ΔZnF1-EGFP, ΔZnF2-EGFP, ΔRRM1-EGFP, ΔRRM2-EGFP, and ΔpNL4-EGFP plasmids for 48 hours. **C** Immunofluorescence was used to detect the effect of the functional domains of MATR3 on its localization in living GO. Green: EGFP, MATR3 forming a fusion protein with EGFP; Blue: Hoechst (nucleus). Scale bar: 10 μm in **B**, 25 μm in **C**.

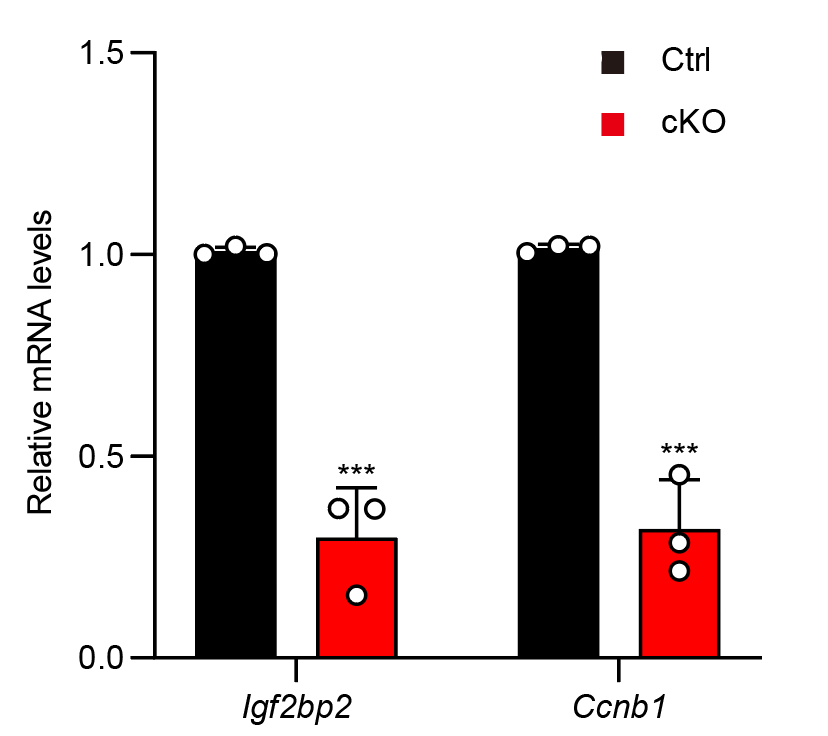

**Figure S9. Targets with high binding intensity and significant expression changes from the LACE-seq data.**

RT-qPCR results showing *Igf2bp2* and *Ccnb1* mRNA level in PD14 oocytes. Data are represented as mean ± SD. ****P* < 0.001.

**Table S1.**

| gRNA | Sequence |
| --- | --- |
| gRNA-1 for *Matr3* ^LoxP/LoxP^ mice | TTAATACGACTCACTATAGGCACCCAAACATCTCACCCTGTTTTAGAGGCTAGAAATAG |
| gRNA-2 for *Matr3* ^LoxP/LoxP^ mice | TTAATACGACTCACTATAGGAAAGCACATTGACTTCAGGGTTTTAGAGGCTAGAAATAG |
| Primers for cloning | Sequence |
| Matr3-SalI-F | ACGCGTCGACTTTCATCCACTGTTCACTAACTCTTCTCTTC |
| Matr3-BamHI-F | GATGGATCCGGTTGGGTGAGATGTTTGGGTGCATCAA |
| Matr3-BamHI-Loxp1-F1 | TGCTATACGAAGTTATCGCCGAGTCCCTTGGATTTAAATGTTTTTAAGCTCAA |
| Matr3-BamHI-Loxp1-F2 | GATGGATCCATAACTTCGTATAATGTATGTATGCTATACGAAGTTATCGCCGAGTCCC |
| Matr3-Nhe-Loxp2-R1 | CATACATTATACGAAGTTATGCGAGAGGGGAAAAATGCTTAAAACAATTCAACA |
| Matr3-Nhe-Loxp2-R2 | ATAGCTAGCATAACTTCGTATAGCATACATTATACGAAGTTATGCGAGAGG |
| Matr3-NheI-F | ATAGCTAGCCTGTCCTGAAGTCAATGTGCTTTCCATTTAGG |
| Matr3-NheI-R | GGAAAAGCGGCCGCAAAGATTTACAGTTCTTTAAGATCATTTTCATT |
| Matr3-screen-F1 | GCCTGCTGTCCTTTGTGGCATTTT |
| Matr3-screen-R1 | ACATTTAAATCCAAGGGACTCGGCG |
| Matr3-screen-R2 | GGAAAGCACATTGACTTCAGGACAGG |
| Matr3-screen-F3 | CACCCAAACATCTCACCCAACCG |
| Matr3-screen-F4 | TGTTTTAAGCATTTTTCCCCTCTCGC |
| Matr3-screen-R3 | TCTGACAATCCTCTTATGTTCCTTTCTTCC |
| Primers for genotyping | Sequence |
| *Matr3*-F | GAAGAAGCCCATCAATGTTTTCACTGCAC |
| *Matr3*-R | AAGGACTGTAGAATCTTCTTCAAGCTGAG |
| *Gdf9*-F | TCTGATGAAGTCAGGAAGAACC |
| *Gdf9*-R | GAGATGTCCTTCACTCTGATTC |

**Table S2.**

| Gene | Forward Primer | Reverse Primer |
| --- | --- | --- |
| *Matr3* | TTGAGAAAAAGAGGGGCGCT | TCGACGACTGTGACTTGCTC |
| *Gdf9* | TCACCTCTACAATACCGTCCGG | GAGCAAGTGTTCCATGGCAGTC |
| *Rdx* | TCAGTGTGACCTTCTCATGCC | AGTCCCATGTCTTGTCTGTGG |
| *β-actin* | CATTGCTGACAGGATGCAGAAGG | TGCTGGAAGGTGGACAGTGAGG |
